# Coordination of humoral immune factors dictates compatibility between *Schistosoma mansoni* and *Biomphalaria glabrata*

**DOI:** 10.1101/767699

**Authors:** Hongyu Li, Jacob R. Hambrook, Emmanuel A. Pila, Abdullah A. Gharamah, Jing Fang, Xinzhong Wu, Patrick C. Hanington

## Abstract

Immune factors in snails of the genus *Biomphalaria* are critical for combating *Schistosoma mansoni*, the predominant cause of human intestinal schistosomiasis. Independently, many of these factors are known to play an important role in, but not fully define, the compatibility between the model snail *B. glabrata,* and *S. mansoni*. Here, we demonstrate association between four, previously characterized humoral immune molecules; *Bg*FREP3, *Bg*TEP1, *Bg*FREP2 and Biomphalysin. We also identify unique immune determinants in the plasma of *S. mansoni*-resistant *B. glabrata* that explain the incompatible phenotype. These factors coordinate to initiate haemocyte-mediated deestruction of *S. mansoni* sporocysts via production of reactive oxygen species. The inclusion of *Bg*FREP2 in a *Bg*FREP3-initiated complex that also includes *Bg*TEP1 almost completely explains resistance to *S. mansoni* in this model. Our study unifies many independent lines of investigation to provide a more comprehensive understanding of the snail immune system in the context of infection by this important human parasite.

## Introduction

Schistosomiasis, a disease caused by parasitic trematodes of the genus *Schistosoma*, is the second-most socioeconomically devastating parasitic disease, with an estimated 252 million people infected worldwide in 2015 (Disease, Injury, & Prevalence, 2016; Mortality & Causes of Death, 2016). Larval digenean trematodes share a common feature in the use of snails (gastropod mollusks) as intermediate hosts for transmission to a vertebrate host (Esch, Barger, & Fellis, 2002). The snail *Biomphalaria glabrata* transmits several species of trematode, including *Schistosoma mansoni*, the predominant causal species of intestinal schistosomiasis (Pila, Gordy, et al., 2016). The molecular interactions between *B. glabrata* and *S. mansoni* have been studied extensively towards better understanding this essential life cycle stage of an important human parasite (Adema, 2015).

Various genetically determined resistance phenotypes exist with respect to *S. mansoni* infection in *B. glabrata* (Richards, 1975a, 1975b). Some *B. glabrata* snails (such as BS-90 strain) naturally resist *S. mansoni* infection. Susceptible *B. glabrata* (such as M-line strain) can develop transient levels of acquired resistance following previous exposure to incompatible or radiation-attenuated parasites with modest cross protection to related digenean species (Lie & Heyneman, 1975; Lie, Heyneman, & Lim, 1975; Sullivan, Richards, Joe, & Heyneman, 1982). Previous research has focused on identifying the mechanisms responsible for determining resistance profiles. Using in vivo and in vitro models of *B. glabrata* snails, it has been demonstrated that the killing of *S. mansoni* larvae is associated with a haemocyte-mediated cytotoxic mechanism, and passive transfer of natural resistance to *S. mansoni* has been successfully accomplished when haemocytes from a susceptible *B. glabrata* strain are incubated in cell-free hemolymph (plasma) from a resistant strain (Bayne, Buckley, & DeWan, 1980a, 1980b; Granath & Yoshino, 1984; Loker & Bayne, 1982). Thus, haemocytes from resistant or susceptible *B. glabrata* strains do not appear to differ a priori in their cytotoxic capabilities, but their response requires activation by some humoral factor(s) (Bayne et al., 1980b; Granath & Yoshino, 1984; Vasquez & Sullivan, 2001), for proper recognition of *S. mansoni* and enhancing haemocyte cytotoxicity (Hahn, Bender, & Bayne, 2001).

Researchers have long sought immune determinants present in resistant *B. glabrata* plasma that specifically activate haemocytes to encapsulate and destroy *S. mansoni* sporocysts. Identified in *B. glabrata*, a large family of soluble lectins termed fibrinogen-related proteins (*Bg*FREPs) attracted widespread interest because of their unique structure (Adema, Hertel, Miller, & Loker, 1997; Gordy, Pila, & Hanington, 2015), capacity for somatic diversification (Zhang, Adema, Kepler, & Loker, 2004), and immune function (Hanington et al., 2010; Hanington & Zhang, 2011). *Bg*FREPs are composed of a C-terminal fibrinogen (FBG) domain and either one or two N-terminal immunoglobulin superfamily (IgSF) domains linked to the FBG domain by an interceding region (ICR) (Hanington et al., 2010; Pila, Li, Hambrook, Wu, & Hanington, 2017). Numerous studies have implicated *Bg*FREP3 as playing a central role in *B. glabrata* resistance to digenetic trematodes (Hanington et al., 2010; Hanington, Forys, & Loker, 2012; Hanington & Zhang, 2011; Lockyer et al., 2012; Lockyer et al., 2008; Pila, Li, et al., 2017). *Bg*FREP3 contains two tandem IgSF domains whereas *Bg*FREP2 has one (Leonard, Adema, Zhang, & Loker, 2001). The expression of *Bg*FREP2 in BS-90 snails is dramatically up-regulated (57-fold) after exposure to *S. mansoni*, whereas M-line snails do not feature such a drastic upregulation, thereby suggesting a role for *Bg*FREP2 in resistance (Hertel, Adema, & Loker, 2005). Interestingly, the time course for heightened *Bg*FREP2 expression overlaps the interval within which sporocysts are encapsulated and killed, and only occurs in BS-90 haemocytes (Dinguirard et al., 2018; Hertel et al., 2005). Furthermore it was reported that *Bg*FREP2 is reactive with polymorphic glycoproteins found on the tegumental surface of larval *S. mansoni* (Mone et al., 2010).

In addition to *Bg*FREPs, other soluble immune effector factors which are involved in both the direct killing of sporocysts and preparation of haemocytes to mount a cell-mediated response have been characterized (Pila, Li, et al., 2017). These factors include *B. glabrata* thioester-containing proteins (*Bg*TEPs) (Bender, Fryer, & Bayne, 1992; Mone et al., 2010; Portet et al., 2018) and Biomphalysin (Galinier et al., 2013). TEPs are key components of invertebrate and vertebrat immune responses, aiding in melanization, opsonization, and killing of invading pathogens (Baxter et al., 2007; Blandin, Marois, & Levashina, 2008; Levashina et al., 2001; Pila, Li, et al., 2017; Portet et al., 2018; Povelones et al., 2013; Yassine & Osta, 2010). *Bg*TEP was first characterized as an alpha macroglobulin proteinase inhibitor, and has recently re-immerged as a molecule of interest due to its association with *Bg*FREP2 and *S. mansoni* polymorphic mucins (*Sm*PoMucs), and the capacity to bind to various invading pathogens (Bender et al., 1992; Mone et al., 2010; Portet et al., 2018). Biomphalysin is a cytolytic protein in *B. glabrata* belonging to the β pore-forming toxin (β-PFT) superfamily (Galinier et al., 2013). Biomphalysin binds to the surface of *S. mansoni* sporocysts in the absence of plasma, while its cytolytic activity is drastically increased when plasma is present, suggesting that other factor(s) within the plasma may mediate the conversion of the oligomeric pre-pore to a functional pore (Galinier et al., 2013). Although the functional mechanisms of these factors are not thoroughly understood, studies suggest that these factors function as key determinants in the final outcome of *S. mansoni* challenge of *B. glabrata* (Galinier et al., 2013; Mone et al., 2010).

While studies have implicated *Bg*FREP3 in resistance to *S. mansoni*, the underlying mechanism of *Bg*FREP3 function; how it binds to *S. mansoni* sporocyst surfaces, and then how recognition is translated into haemocyte engagement, activation, and ultimately parasite encapsulation, is still unknown (Hanington et al., 2010). Here, we report an association between *Bg*FREP3, *Bg*TEP1, Biomphalysin (UniProtKB/TrEMBL: A0A182YTN9 and A0A182YTZ4) and *Bg*FREP2 (UniProtKB/TrEMBL: A0A2C9L9F5). *Bg*FREP3 associates with *Bg*TEP1 and Biomphalysin in both M-line and BS-90 strains, but only in BS-90 strain uniquely interacts with *Bg*FREP2 and other versions of *Bg*FREP3 (a variant of *Bg*FREP3.3), providing us with an important insight into why BS-90 strain is refractory to *S. mansoni* infection. In this study, we demonstrate that *Bg*FREP3 binds to *S. mansoni* sporocysts without any other soluble plasma factors, yet *Bg*FREP2 relies on *Bg*TEP1. However, *Bg*FREP3 still interacts with *Bg*TEP1 to form unique immune complexes, which significantly enhance the ability of haemocytes and plasma from *S. mansoni*-susceptible *B. glabrata* (M-line) to kill *S. mansoni* sporocysts. A more striking finding is that the combination of *Bg*FREP3, *Bg*TEP1 and *Bg*FREP2 renders M-line haemocytes capable of killing *S. mansoni* sporocysts at nearly the same level as *S. mansoni*-resistant BS-90 haemocytes. Reactive oxygen species (ROS) is shown to play an important role during this haemocyte-mediated killing of *S. mansoni* sporocysts. These results provide insight into how the numerous previously characterized immune factors known to be important in the anti-*S. mansoni* immune response to *B. glabrata* are acting in concert to defend the snail host.

## Results

### *Bg*FREP3 interacts with *Bg*TEP1, Biomphalysin, *Bg*FREP2 and other *Bg*FREP3 variants in snail plasma

To identify the factors in snail plasma which interact with *Bg*FREP3, we produced recombinant *Bg*FREP3 (r*Bg*FREP3, GenBank: AAK28656.1) using an insect expression system (Fig. 1A and Figure 1 – figure supplement 1) and performed a series of pull-down experiments. We initiated our investigations with *Bg*FREP3 because we observed that it is expressed at greater abundance in BS-90 compared with M-line snails. This observation was confirmed using Western-blot that showed that the hemolymph of BS-90 snails contain more *Bg*FREP3 than does M-line snail hemolymph (Fig. 1B). Among the proteins identified using liquid chromatography-tandem mass spectrometry (LC-MS/MS) in the eluent of these pull-down experiments was *Bg*TEP1 (Fig. 1C). We performed four r*Bg*FREP3 pull-down experiments, each yielded a band at approximately 200-kDa, which corresponded to the full-length *Bg*TEP1, which was identified from both M-line and BS-90 strains (Fig. 1C). In three instances, we cut the bands and sent for mass spectrometry identification, two of which detected the *Bg*TEP1 pulled down by r*Bg*FREP3 in M-line plasma, while the third detected *Bg*TEP1 in both M-line and BS-90 strains (Fig. 1D). From the LC-MS/MS analysis we obtained a total of 17 unique peptides, 16 peptides were detected from M-line strain, 4 peptides were from BS-90 strain, and 3 peptides were present in both BS-90 and M-line strains (Table. S2 and Fig. 1E). While all peptides mapped to *Bg*TEP, there is uncertainty as to which specific *Bg*TEP proteins are present. Twelve peptides mapped to *Bg*TEP-ACL00841.1; 15 peptides mapped to *Bg*TEP1.1-ADE45332.1, *Bg*TEP1.2-ADE45339.1, *Bg*TEP1.3-ADE45340.1, *Bg*TEP1 1.4-ADE45341.1 and *Bg*TEP1.5-ADE45333.1, however, there was overlap between some of these peptides, which mapped to all known *Bg*TEP sequences (Figure 1 – figure supplement 2). The identified *Bg*TEP1 showed a relatively large difference from *Bg*TEP (ACL00841.1), but could not be distinguished from other *Bg*TEP1 sequences (Figure 1 – figure supplement 2), so we termed it *Bg*TEP1.

**Figure 1.**
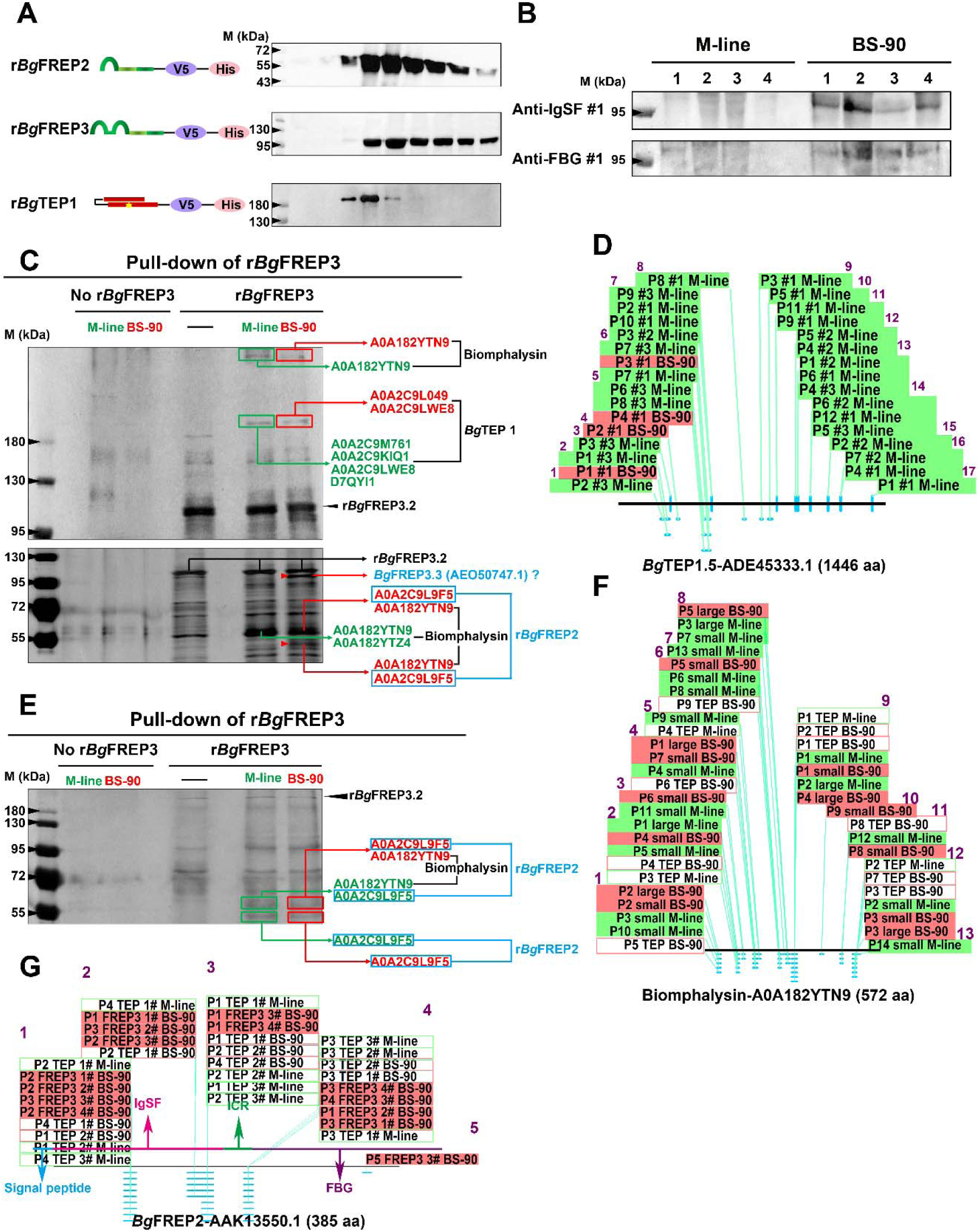
Pull-down experiments of r*Bg*FREP3 and r*Bg*TEP1. **A:** Schematic diagram and Western blot analysis of recombinant proteins, the ends of which are coupled to the tags V5 and ploy-His for protein immunoassay and purification operations. **B:** Western blot analysis of *Bg*FREP3 background expression in hemolymph of M-line and BS-90 snails. The hemolymph of four snails from each M-line and BS-90 strain was used for detection, and anti-IgSF #1 and anti-FBG #1 antibodies were used as primary antibodies. **C:** Pull-down experiments with r*Bg*FREP3. The eluate was collected and separated on 6% (upper panel) and 10% (lower panal) SDS-PAGE before silver staining. **E:** Pull-down experiments with r*Bg*TEP1. The eluate was separated on 10% SDS-PAGE. In the pull-down experiment, cell-free plasma from susceptible (green) or resistant (red) *B. glabrata* snails was incubated with Sf9 cell lysates expressing r*Bg*FREP3 or concentrated medium containing r*Bg*TEP1 together (interactome experiments, last two lanes) or alone (controls, first three lanes). Bands that differ between control and interactome experiments were cut, proteins submitted to tryptic digest and analyzed by LC-MS/MS for identification. In interactome experiments, bands that differ between M-line and BS-90 were marked with red arrows. The results of mass spectrometry analysis are shown in the figure. **D, F and G:** Distribution of peptides identified by LC-MS/MS on *Bg*TEP1.5 (ADE45333.1), Biomphalysin (A0A182YTN9) and *Bg*FREP2 (AAK13550.1). Green background: from M-line strain in the r*Bg*FREP3-pull-down; red background: from BS-90 strain in the r*Bg*FREP3-pull-down; green box: from M-line strain in the r*Bg*TEP1-pull-down; red box: from BS-90 strain in the r*Bg*TEP1-pull-down.

The full-length sequence of *Bg*TEP1 generally contains 1445 or 1446 amino acids (Figure 1 – figure supplement 2). The peptides identified by LC-MS/MS cover 16.3% of the full-length sequence (Figure 1 – figure supplement 2). The identified peptides are distributed over the full-length *Bg*TEP1, with the peptides identified from the BS-90 strain being concentrated in the N-terminal region (Fig. 1D). Background transcript abundance of *Bg*TEP1 detected by qRT-PCR in M-line and BS-90 snails, suggest that there is no significant difference (P > 0.05) in baseline *Bg*TEP1 expression between the two strains (Figure 1 – figure supplement 3). These results indicate that the interaction of *Bg*FREP3 with *Bg*TEP1 doesn’t require the involvement of any parasite components.

Recombinant *Bg*TEP1 (r*Bg*TEP1, GenBank: ADE45332.1) was also used to perform a series of pull-down experiments (Fig. 1A and E). Biomphalysin (UniProtKB/TrEMBL: A0A182YTN9 and A0A182YTZ4) was identified in both M-line and BS-90 plasma from r*Bg*TEP1 pull-down experiments and was also found to be present in the pull-down studies using *Bg*FREP3 (Fig. 1C and E). Analysis suggests that this Biomphalysin (A0A182YTZ4) was most similar (97% amino acid identity) to the Biomphalysin previously reported by Galinier and Portela *et al*. (Galinier et al., 2013) (GenBank: KC012466) (Figure 1 – figure supplement 4). The 13 peptides identified by LC-MS/MS covered 28.3% of Biomphalysin (A0A182YTN9) amino acid sequence (Fig. 1F, Table S2 and Figure 1 – figure supplement 4). These peptides were evenly distributed throughout the Biomphalysin protein (Fig. 1F). The theoretical molecular weight of Biomphalysin (A0A182YTN9 and A0A182YTZ4) is ∼64-kDa. The Biomphalysin we identified via protein pull-down emerged at three different molecular weights, one at > 400-kDa (Fig. 1C), which likely represents the heptamer formed by Biomphalysin (Galinier et al., 2013), the second site at roughly 60-kDa (Fig. 1C and E), which is in agreement with the theoretical molecular weight, and the third at around 55-kDa (Fig. 1C), which may represent the size after proteolysis (Galinier et al., 2013).

Comparing r*Bg*FREP3 pull-down results between M-line and BS-90 *B. glabrata* identified 2 unique proteins in the BS-90 lane. One of these proteins is *Bg*FREP2 (UniProtKB/TrEMBL: A0A2C9L9F5) (Fig. 1C and E). Five peptides were identified by LC-MS/MS analysis and covered 16.6% of *Bg*FREP2 (GenBank: AAK13550.1) amino acid sequence and were located throughout the IgSF and FBG of the protein (Fig. 1G and Table S2 and Figure 1 – figure supplement 5). Three of the five identified peptides specifically belonged to *Bg*FREP2 (AAK13550.1), so the identified *Bg*FREP2 appears to represent a specific *Bg*FREP2 variant that is distinguished from other *Bg*FREP members that have been published (Figure 1 – figure supplement 5).

Only *Bg*FREP2 was identified, at ∼55-kDa, from the BS-90 strain plasma when r*Bg*FREP3 was used as the bait protein in pull-down experiments (Fig. 1C). This contrasts with our other data in which *Bg*FREP2 was identified from plasma of both M-line and BS-90 strains when r*Bg*TEP1was used as bait in pull-down experiments (Fig. 1E). *Bg*FREP2 was identified by LC-MS/MS from two distinct molecular weight bands in the r*Bg*TEP1 study, one band, at ∼60 kDa also included peptides that mapped to Biomphalysin, the other band, identified at ∼55-kDa was contained peptides for only *Bg*FREP2 (Fig. 1C and E). Thus, all of the *Bg*FREP2 peptides were identified from protein bands that were of greater molecular weight than the predicted *Bg*FREP2, which is 43.8-kDa. However, this is consistent with the observed molecular weight of *Bg*FREP2 by Moné *et al*. (Mone et al., 2010), and could be explained by post-translational modifications such as glycosylation (Adema et al., 1997; Mone et al., 2010; Zhang, Zeng, & Loker, 2008).

Another unique protein identified from the BS-90 lane was a variant of *Bg*FREP3. Of the 9 peptides identified by LC-MS/MS, 7 specifically matched *Bg*FREP3.3 (Genbank: AEO50747.1), the other two matched other members of *Bg*FREP3 family (*Bg*FREP3.3-AAO59915.1, *Bg*FREP3.2-AEO50746.1, *Bg*FREP3.2-AAK28656.1 and *Bg*MFREP3-AAK13548.1), suggesting that this protein is likely a FREP3 variant (Figure 1 – figure supplement 6 and Table S2). The 7 peptides that specifically belonged to *Bg*FREP3.3 (AEO50747.1) covered 15.6% of the amino acid sequence length and mainly concentrated in the interceding region (ICR) (5 peptides), one peptide in each of the IgSF2 domain and FBG domain of *Bg*FREP3.3 were also identified (Figure 1 – figure supplement 6).

### *Bg*FREP3 independently recognizes and binds to primary sporocysts of *S. mansoni*

Our data suggests a close and complex interaction between *Bg*FREP3, *Bg*TEP1, Biomphalysin and *Bg*FREP2. These factors have each independently been reported to be involved in the recognition of *S. mansoni* sporocysts (Adema et al., 1997; Galinier et al., 2013; Hanington et al., 2010; Hanington et al., 2012; Mone et al., 2010; Portet et al., 2018; X. J. Wu et al., 2017). However, only Biomphalysin is known to directly bind to the surface of *S. mansoni* sporocysts without the aid of any other plasma factors (Galinier et al., 2013). The mechanism by which *Bg*FREP3, *Bg*TEP1 and *Bg*FREP2 bind to primary sporocysts of *S. mansoni* is still not clear. To explore this issue, we produced r*Bg*FREP3, r*Bg*TEP1 and r*Bg*FREP2 (Fig. 1A) to observe whether they associate with primary sporocysts of *S. mansoni*. Immunocytochemistry clearly shows that r*Bg*FREP3 is able to interact with *S. mansoni* sporocysts independently, while r*Bg*TEP1 and r*Bg*FREP2 alone do not have such capabilities (Fig. 2). However, incubation of sporocysts with pre-combined r*Bg*FREP2 and r*Bg*TEP1 yields a signal, suggesting that these two factors require complex formation prior to being able to associate with *S. mansoni* sporocysts (Fig. 2).

**Figure 2.**
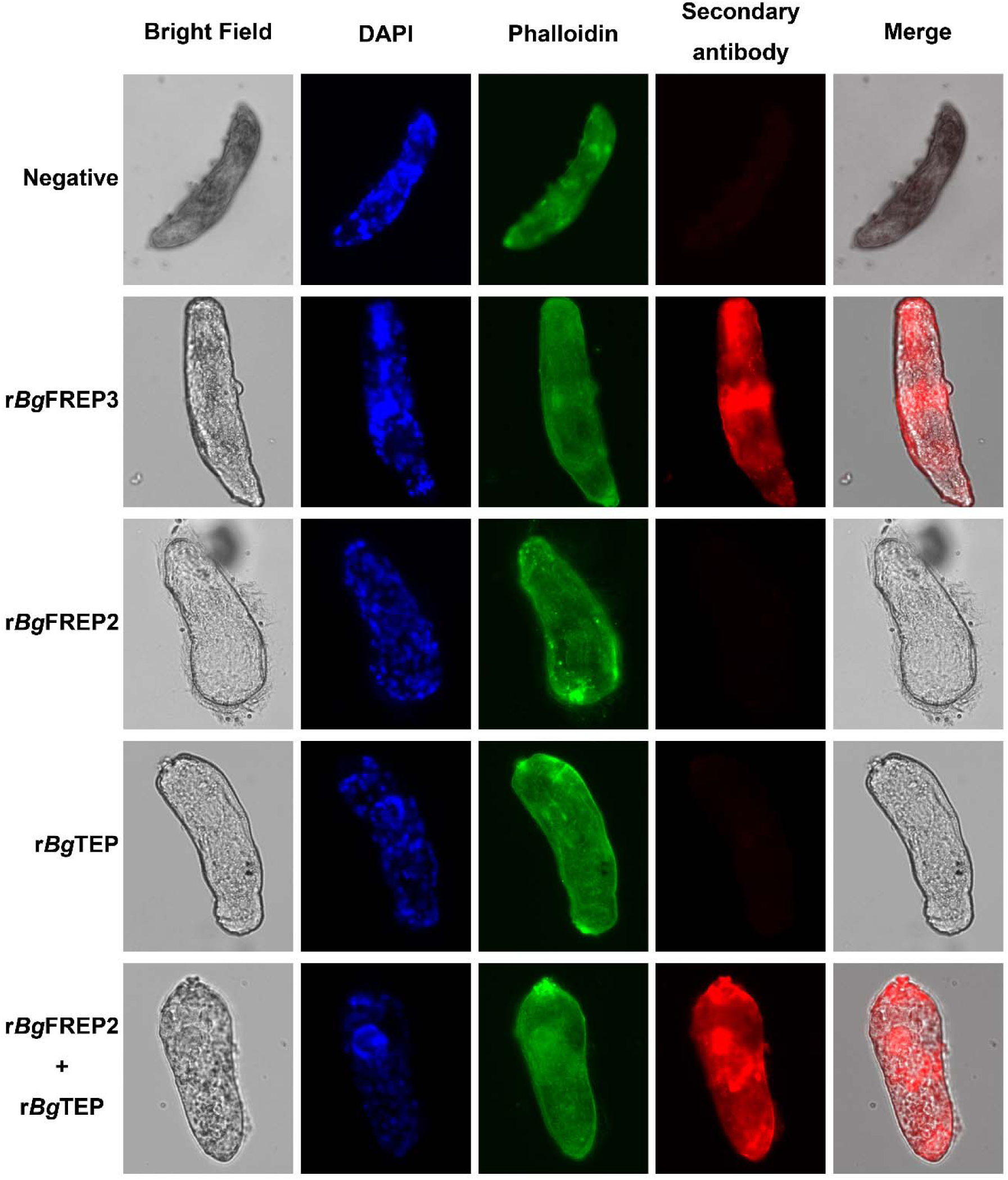
Immunofluorescence analysis of interaction between r*Bg*FREP3, r*Bg*FREP2, and r*Bg*TEP1 with *S. mansoni* sporocysts. Sporocysts were incubated with vehicle or recombinant proteins (r*Bg*FREP3, r*Bg*FREP2, and r*Bg*TEP1 alone) and combination (r*Bg*FREP2 plus r*Bg*TEP1) and immunostained using anti-V5 primary IgG and Alexa Fluor 633 goat anti-mouse secondary antibody. The sporocysts were stained with DAPI (for nucleus, blue), phalloidin (for F-actin labeling, green) and fluorescent secondary antibody (for recombinant proteins, red).

### *Bg*FREP3 and *Bg*TEP1 form an immune complex

To further investigate whether *Bg*FREP3 and *Bg*TEP1 form a complex, we carried out a set of immunoblotting experiments with recombinant proteins. As expected, r*Bg*FREP3 did form a complex with r*Bg*TEP1 (Fig. 3A). The complex appeared in a region of high molecular weight (> 460-kDa), and the signal decreased gradually with the decrease of r*Bg*TEP1:r*Bg*FREP3 ratio in a dose-dependent manner (Fig. 3A). As shown in Fig. 3A, r*Bg*FREP3 exists as a high molecular weight multimer in a non-denatured state. Since the molecular weights of the r*Bg*FREP3 multimer and the r*Bg*FREP3-r*Bg*TEP1 complex were both > 460-kDa, it was difficult to distinguish them using Western bolt. To confirm the interaction between r*Bg*FREP3 and r*Bg*TEP1, a far-Western blotting approach, in which decreasing amount of r*Bg*TEP1 was imprinted on the membrane, and then detected by using biotinylated r*Bg*FREP3 as primary recognition agent was performed. A band was detected in the lane with the largest amount of r*Bg*TEP1, confirming the interaction between r*Bg*FREP3 and r*Bg*TEP1 (Fig. 3B).

**Figure 3.**
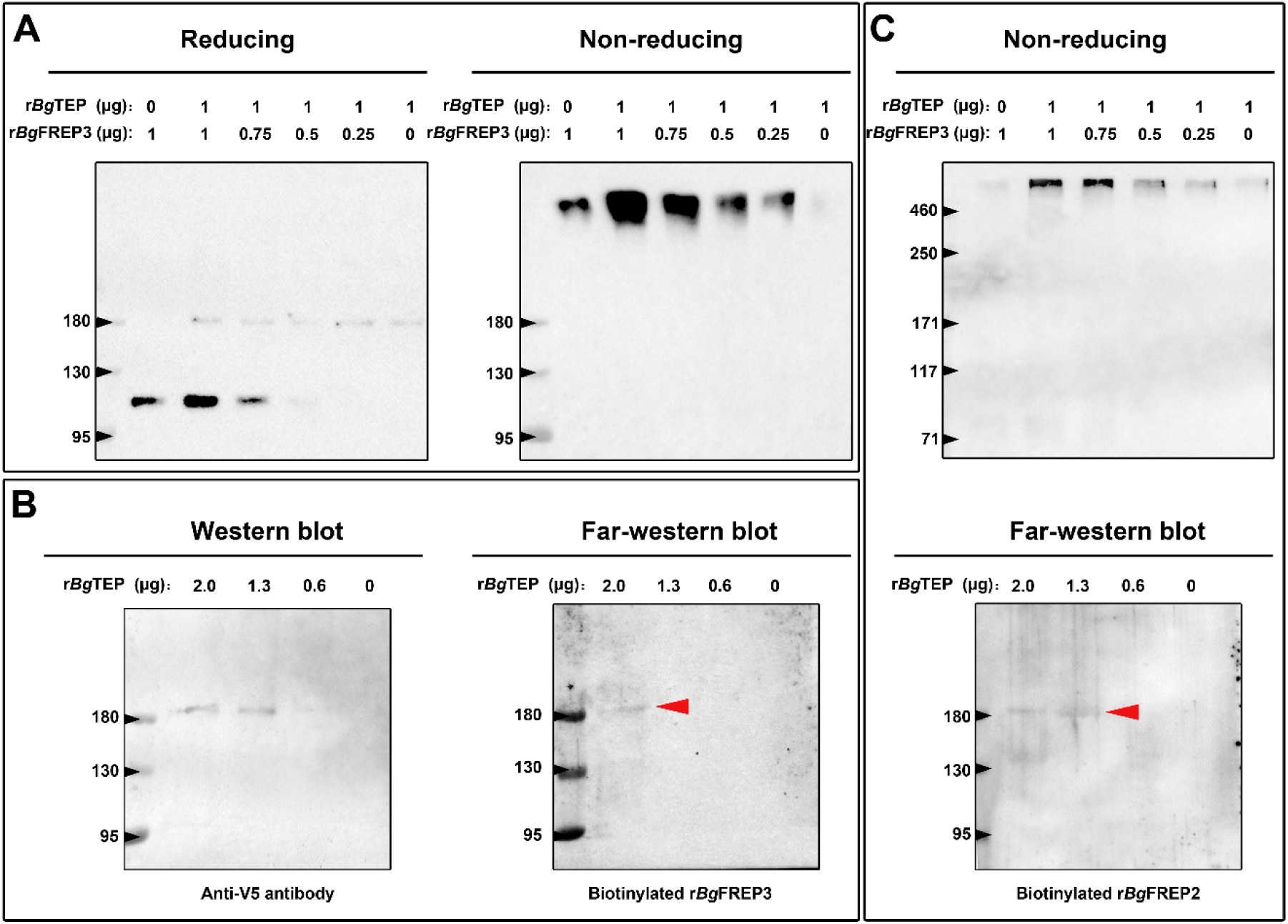
Immunoblot analysis of the interaction between r*Bg*TEP1 and r*Bg*FREP3 and r*Bg*FREP2. **A:** Purified r*Bg*TEP1 was incubated with r*Bg*FREP3 in the indicated ratios (wt/wt in µg) for 2 h at room temperature. After the incubation, Western blot analysis was carried out under denaturing (heating with reducing agent, the right) and non-denaturing (no heating, no reducing agent, the left) conditions, respectively. **B:** Decreasing amount of r*Bg*TEP1 was blotted onto the membrane and then subjected to standard Western blot (the left) and far-Western blot (the right) analysis. **C:** The interaction between r*Bg*FREP2 and r*Bg*TEP1. Western blot analysis under non-denaturing (the upper) and far-Western blot analysis (the below) was carried out, respectively. In the far-Western blot, the biotinylated r*Bg*FREP3 and r*Bg*FREP2 acts as a primary antibody and the secondary antibody is a streptavidin-HRP; the red arrow indicates the band that symbolizes the interaction between *Bg*TEP1 and r*Bg*FREP3 or r*Bg*FREP2.

*Bg*FREP2 and *Bg*TEP1 have been shown to form immune complexes with *Sm*PoMucs derived from *S. mansoni* (Mone et al., 2010), suggesting that *Bg*FREP2 and *Bg*TEP1 associate with each other in snail plasma. To test this assumption, we performed a set of immunoblot assays using r*Bg*FREP2 and r*Bg*TEP1. The results of Western blot under non-denaturing conditions showed that a protein at > 460 kDa after mixing r*Bg*FREP2 and r*Bg*TEP1. This protein decreased in intensity on the Western blot in a dose dependent manner along with the decrease of the r*Bg*TEP1:r*Bg*FREP2 ratio (Fig. 3C upper panel). However, r*Bg*FREP2 does not exist as a multimer under non-denaturing conditions (Fig. 3C upper panel), which is consistent with the findings of Zhang *et al*. that *Bg*FREP2 did not form a multimer in snail plasma (Zhang et al., 2008). In addition, we further verified the interaction between r*Bg*FREP2 and r*Bg*TEP1 by Far-Western blot (Fig. 3C lower panel).

### Association of *Bg*FREP3 and *Bg*TEP1 facilitates recognition and killing of *S. mansoni* sporocysts

Since *Bg*FREP3 does not rely on *Bg*TEP1 to recognize pathogens, but does associate with *Bg*TEP1, we hypothesize that *Bg*TEP1 plays a role in the downstream immune response triggered by *Bg*FREP3 recognition of *S. mansoni* sporocysts. We treated primary sporocysts with haemocytes or plasma from BS-90 and M-line *B. glabrata* snails with r*Bg*FREP3, r*Bg*TEP1 and a combination of the two in vitro.

Plasma (hemoglobin-low/free) from the resistant BS-90 strain of *B. glabrata* snails is naturally able to kill 67% (SEM 3.7%; n=10) of *S. mansoni* sporocysts by 48 h post incubation, while plasma from the susceptible M-line strain, which kills 20% (SEM 4.5%; n=10), is not significantly more effective than the medium control in which 13% (SEM 2.1%; n=10) were killed at 48 h (Fig. 4A). The addition of r*Bg*FREP3 (36% [SEM 5.0%; n=10]) and r*Bg*TEP1 (32% [SEM 3.3%; n=10]) independently did not significantly enhanced the ability of M-line plasma to kill *S. mansoni* sporocysts by 48 h (Fig. 4A). However, the combined addition of r*Bg*FREP3 and r*Bg*TEP1 did significant increase M-line-plasma-mediated killing of *S. mansoni* sporocysts by 48 h post incubation (56% [SEM 4.8%; n=10]) compared to the application of either recombinant protein alone. The combined addition of r*Bg*FREP3 and r*Bg*TEP1 rendered the capacity of M-line plasma able to kill *S. mansoni* sporocysts statistically insignificant from BS-90 plasma (P > 0.05) (Fig. 4A). Recombinant proteins, in the absence of *B. glabrata* plasma, did not display any capacity to kill *S. mansoni* sporocysts (Figure 4 – figure supplement 1A). This suggests that r*Bg*FREP3 and r*Bg*TEP1 activate a factor in plasma to confer the ability to kill sporocysts.

**Figure 4.**
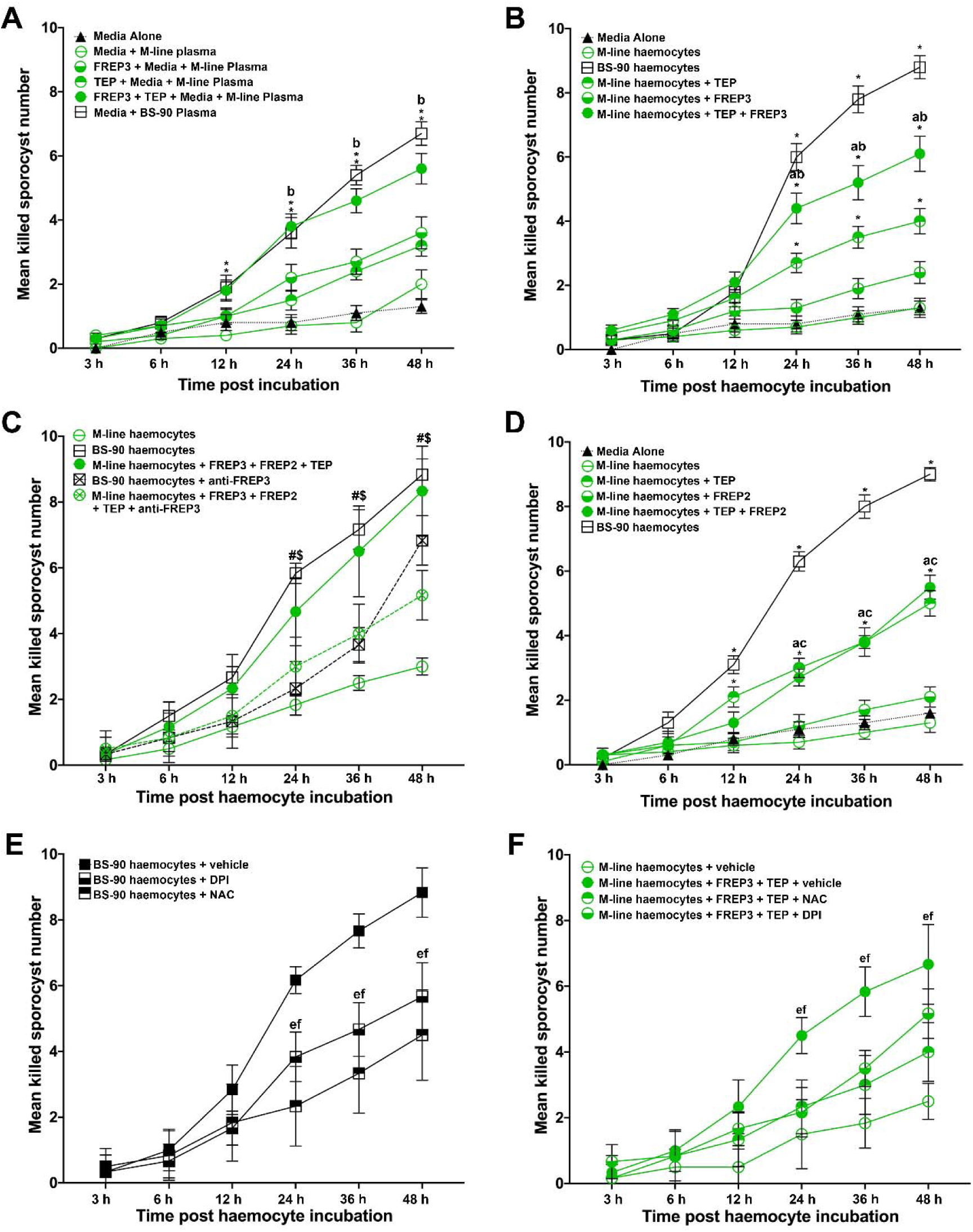
Association of r*Bg*FREP3, r*Bg*TEP1 and r*Bg*FREP2 facilitates M-line snails to kill *S. mansoni* sporocytes. Effects of r*Bg*FREP3, r*Bg*TEP1 and r*Bg*FREP3-r*Bg*TEP1 complex on killing of *S. mansoni* sporocysts by M-line plasma (**A**) and haemocytes (**B**). **C:** Before being exposed to sporocysts, M-line haemocytes were pre-incubated with r*Bg*FREP3-r*Bg*TEP1-r*Bg*FREP2 combination in presence or in absence of anti-*Bg*FREP3 antibodies (abrogation treatment). BS-90 haemocytes were incubated with anti-*Bg*FREP3 antibodies. **D:** The effect of r*Bg*FREP2, r*Bg*TEP1 and r*Bg*FREP2-r*Bg*TEP1 complex on the destruction of sporocysts by M-line haemocytes. **E and F:** The role of ROS in killing of *S. mansoni* sporocysts by haemocytes from *B. glabrata* snails. **E:** The addition of ROS inhibitors NAC and DPI abolished the ability of BS-90 haemocytes to destroy sporocysts. **F:** Killing sporocysts of M-line haemocytes rendered by r*Bg*FREP3-r*Bg*TEP1 complex was annulled by pre-incubation with ROS inhibitors NAC and DPI. Statistically significant difference symbols in **A**, **B** and **D**: asterisk (*) represents comparison with M-line haemocytes or plasma, * P < 0.05, ** P < 0.01; a represents comparison with BS-90 haemocytes or plasma, P < 0.05; b represents comparison with both single treatment (r*Bg*TEP1/r*Bg*FREP3 or r*Bg*FREP2), P < 0.05; c represents comparison with only *Bg*FREPs treatment, P < 0.05; bars represent SEM, n=10. Statistically significant difference symbols in **C**: # represents comparison BS-90 haemocytes with BS-90 haemocytes+anti-r*Bg*FREP3 antibodies, P < 0.05; $ represents comparison M-line haemocytes+r*Bg*FREP3+r*Bg*FREP2+r*Bg*TEP1 with M-line haemocytes+r*Bg*FREP3+r*Bg*FREP2+r*Bg*TEP1+anti-r*Bg*FREP3 antibodies, P < 0.05; bars represent SEM, n=10. Statistically significant difference symbols in **E** and **F:** e represents comparison NAC treatment with BS-90 haemocytes or M-line haemocytes+r*Bg*FREP3+r*Bg*TEP1, P < 0.05; f represents comparison DPI treatment with BS-90 haemocytes or M-line haemocytes+r*Bg*FREP3+r*Bg*TEP1, P < 0.05; bars represent SEM, n=6.

Haemocytes derived from BS-90 strain *B. glabrata* are innately capable of destroying *S. mansoni* sporocysts, killing 88% (SEM 3.6%; n=10) of sporocysts after 48 h of incubation in vitro (Fig. 4B). The basal ability of M-line haemocytes to kill sporocysts (13% [SEM 3%; n=10]) is not significantly different from the medium-alone control (13% [SEM 2.1%; n=10]), and is only slightly increased by adding r*Bg*FREP3 (24% [SEM 3.4%; n=10]) at 48 h (Fig. 4B). Addition of r*Bg*TEP1 did significantly increase sporocyst killing by M-line haemocytes to 40% (SEM 3.9%; n=10) (Fig. 4B), however, this killing response was significantly enhanced to 61% (SEM 5.5%; n=10) when r*Bg*FREP3 and r*Bg*TEP1 were incubated with M-line haemocytes together (Fig. 4B). However, the combination of r*Bg*FREP3 and r*Bg*TEP1 did not convey a killing capacity to M-line snails that paralleled BS-90 haemocytes (Fig. 4B). This suggests that there is yet another factor(s) affecting this process, causing haemocytes from BS-90 and M-line to differ in their ability to destroy sporocysts. In our r*Bg*FREP3 pull-down experiments, *Bg*FREP2 was one of the two proteins uniquely identified as associating with r*Bg*FREP3 in the plasma of BS-90 *B. glabrata* (Fig. 1C). Incubation of r*Bg*FREP2 in combination with r*Bg*FREP3 and r*Bg*TEP1 enabled M-line haemocytes to kill *S. mansoni* sporocysts (83% [SEM 4.8%; n=6]) to statistically indistinguishable (p > 0.05) levels as BS-90 haemocytes (88% [SEM 4.7%; n=6])) (Fig. 4C). To verify whether the r*Bg*FREP2-r*Bg*TEP1 complex dramatically enhanced the ability of M-line haemocytes to killing *S. mansoni* sporocysts above either factor independently, as does the r*Bg*FREP3-r*Bg*TEP1 complex, a similar experiment was performed, showing that r*Bg*FREP2-r*Bg*TEP1 did not play such a role (Fig. 4D). Independent treatments of r*Bg*FREP2 (21% [SEM 3.1%; n=10]) and r*Bg*TEP1 served controls (Figure 4 – figure supplement 1B), demonstrating that *Bg*FREP2 on alone does not facilitate sporocyst killing above control, whereas r*Bg*TEP1 killing (50% [SEM 3.9%; n=10]) is indistinguishable (P > 0.05) from r*Bg*FREP2 and r*Bg*TEP1 combined (55% [SEM 3.7%; n=10]) (Fig. 4D).

These results made us speculate that *Bg*FREP3 was the determining factor controlling sporocyst engagement and killing. Addition of three anti-r*Bg*FREP3 antibodies to the mixture of r*Bg*FREP3, r*Bg*TEP1 and r*Bg*FREP2 significantly abrogates the increase in the ability of M-line haemocytes to kill sporocysts (51% [SEM 3.1%; n=6] compared to 83% [SEM 4.8%; n=6]) (Fig. 4C). Pre-incubation of BS-90 haemocytes with the anti-r*Bg*FREP3 antibodies also significantly reduced their ability to destroy sporocysts (68% [SEM 3.1%; n=6] compared to 88% [SEM 4.7%; n=6]) (Fig. 4C). Combined pre-immunized IgG purified from serum from the same rabbits from which the antibodies were produced was used as a negative control, demonstrating that it did not significantly affect the killing *S. mansoni* sporocysts by M-line (51% [SEM 3%; n=6]) and BS-90 haemocytes (90% [SEM 3.7%; n=6]) (Figure 4 – figure supplement 1C). These results suggest that r*Bg*FREP3 plays a central role in initiating snail haemocyte-mediated anti-*S. mansoni* immune responses.

Finally, we hypothesize that haemocyte-mediated killing likely involves the production of ROS, which have been shown to be crucial to the haemocyte-mediated clearance of *S. mansoni* sporocysts (Hahn, Bender, & Bayne, 2000; Hahn et al., 2001). To investigate the role of ROS in the combined ability of r*Bg*FREP3 and r*Bg*TEP1 to enhance the ability of M-line haemocytes to kill *S. mansoni* sporocysts, ROS inhibitors N-acetyl-L-cysteine (NAC) and Diphenyleneiodonium (DPI) were applied to the sporocyst killing assays. Co-incubation of BS-90 haemocytes with NAC and DPI significantly reduced their ability to kill *S. mansoni* sporocysts (45% [SEM 5.6%; n=6] and 57% [SEM 4.2%; n=6] respectively) compared to BS-90 haemocytes in the vehicle control (89% [SEM 3.1%; n=6]) by 48 h post incubation (Fig. 4E). The ability of M-line haemocytes primed by the r*Bg*FREP3-r*Bg*TEP1 complex to destroy sporocysts (67% [SEM 4.9%; n=6]) was also significantly abrogated by the addition of NAC (40% [SEM 3.7%; n=6]) or DPI (52% [SEM 3.1%; n=6]) (Fig. 4F).

## Discussion

In 1984, Yoshino *et al*. successfully achieved passive transfer of resistance to *S. mansoni* from a refractory *B. glabrata* strain to susceptible snails via injection of cell-free hemolymph (plasma) (Granath & Yoshino, 1984). Similar passive transfer of resistance had previously been shown using an in vitro model (Bayne et al., 1980a, 1980b; Loker & Bayne, 1982). Although the oxidation of haemoglobin, which is the main protein component (97% of total protein) in *B. glabrata* plasma was later found to be toxic to *S. mansoni* sporocysts in long-term cultures (Bender, Bixler, Lerner, & Bayne, 2002), and thus likely explained the passive transfer study results, further studies suggest that specific anti-parasitic factors must be present at or above a certain threshold level in *S. mansoni*-resistant snail plasma to stimulate a haemocyte-mediated destruction of sporocysts within 24–48 h post-infection (Dinguirard et al., 2018). Although many important immune factors have been identified from *B. glabrata* plasma/haemocytes, such as *Bg*FREPs (Adema et al., 1997), *Bg*TEP1 (Mone et al., 2010), Biomphalysin (Galinier et al., 2013), Toll-like receptors (*Bg*TLRs) (Pila, Tarrabain, Kabore, & Hanington, 2016), granulin (*Bg*GRN) (Pila, Gordy, et al., 2016) and macrophage migration inhibitory factor (*Bg*MIF) (Baeza Garcia et al., 2010), it is still not known which factors ultimately underpin Biomphalaria immunity *S. mansoni*. Here, we provide insight into the broader interactions that take place between *B. glabrata* immune factors that leads to development of an *S. mansoni*-resistant phenotype.

Previous studies have shown that *Bg*FREPs are capable of binding to digenean trematode sporocysts and are up-regulated in response to larval infection (Adema et al., 1997; X. J. Wu et al., 2017; Zhang, Leonard, Adema, & Loker, 2001). *Bg*FREP3 has been shown to be a central part of the anti-*S. mansoni* immune response (Hanington et al., 2010). *Bg*FREP3 is up-regulated based on three characteristics (size, strain, and acquired resistance) of resistance of *B. glabrata* to infection with *S. mansoni* or *Echinostoma paraensei* (Hanington et al., 2010). Knock-down of *Bg*FREP3 in resistant snails using siRNA-mediated interference altered the phenotype of resistant *B. glabrata* by increasing susceptibility to *E. paraensei* infection (Adema, 2015; Hanington et al., 2010). Nevertheless, we have little knowledge of the underlying mechanism by which *Bg*FREP3 interacts with other snail plasma factors in order to signal and trigger an immune response after recognizing the pathogen. In order to explore this mechanism, we conducted a series of pull-down experiments that allowed us to discover the interaction between *Bg*TEP1, Biomphalysin and *Bg*FREP2 with *Bg*FREP3 (Fig. 1).

Proteomic analysis of *B. glabrata* plasma revealed that *Bg*FREP3, *Bg*TEP1 and *Bg*FREP2 display affinity for *S. mansoni* sporocyst tegumental membrane proteins (Portet et al., 2018; X. J. Wu et al., 2017). *Bg*TEP1 was previously reported to play a role in *Bg*FREP2 recognition of *S. mansoni* sporocyst *Sm*PoMucs, (Mone et al., 2010; Portet et al., 2018). Our immunofluorescence results illustrate that *Bg*FREP3 independently recognizes *S. mansoni* sporocysts, whereas *Bg*FREP2 must be involved with the participation of *Bg*TEP1 (Fig. 2). We speculate that this may be due to their structural differences: *Bg*FREP3 possesses two IgSF domains, yet *Bg*FREP2 has only one; *Bg*FREP3 exists as a multimer in its natural state (Fig. 3), whereas *Bg*FREP2 does not (Zhang et al., 2008). Alternatively, it is possible that the targets they recognize are different, so the immunoaffinity reactions that occur are also fundamentally different. A recent report suggests that *Bg*FREPs are likely to form multimers through the coiled-coil region of the ICR. However, this has not been supported by experimental evidence and does not completely rule out the possibility that *Bg*FREP multimer formation is mediated by the FBG or IgSF domain (Gorbushin, 2019).

*Bg*TEP1 and *Bg*FREP3 form a complex without any *S. mansoni* molecules present (Fig. 3). Since *Bg*FREP3 does not require *Bg*TEP1 to recognize pathogens, *Bg*TEP1 is more likely to play a role in the downstream immune response initiated by *Bg*FREP3. Following the addition of r*Bg*FREP3 and r*Bg*TEP1, plasma demonstrated an increased capacity to kill sporocysts compared to controls (Fig. 4A). Because the *B. glabrata* plasma used in these tests had haemoglobin removed, the toxicity of hemoglobin oxidation to *S. mansoni* sporocysts was ruled out. These studies also demonstrated that recombinant *Bg*FREP3 and *Bg*TEP1 alone were not able to increase sporocyst killing, but showed that the addition of cell-free plasma with both recombinants was required, thereby implying the necessity of another factor. Based on our pull-down experiments, we think that this factor could be a Biomphalysin. Biomphalysin displays similarities to members of the β-PFT superfamily (Howard & Buckley, 1985; MacKenzie, Hirama, & Buckley, 1999; Wilmsen, Leonard, Tichelaar, Buckley, & Pattus, 1992; Xu, Wang, Yu, & Sun, 2014), and has cytolytic activity mediated by a plasma factor(s) that remains to be identified (Galinier et al., 2013). Our results suggest a scenario in which *Bg*FREP3 and/or *Bg*TEP1 could be the factors associating with Biomphalysin to promote plasma-mediated sporocyst killing. *Bg*FREP3 and/or *Bg*TEP1 might activate Biomphalysin in an unknown manner, assist in the oligomerization of Biomphalysin while it forms its heptameric channel, or mediate the conversion of the oligomeric prepore to a functional pore. Further investigation is warranted to illustrate the role of *Bg*FREP3 and *Bg*TEP1 in the activation of Biomphalysin.

Within 2–3 h of *S. mansoni* entry into *B. glabrata*, haemocytes surround the developing sporocysts forming multi-layered cellular capsules that kill encapsulated larvae within 24–48 h post-infection (Dinguirard et al., 2018; Loker & Bayne, 1982). This occurs in both susceptible and resistant snail strains and has been replicated in our in vitro functional studies, where only resistant snails can eventually destroy the sporocysts without external assistance. Dinguirard *et al*. conducted a comparative proteomic analysis on the responses of haemocytes participating in *S. mansoni* sporocyst encapsulation reactions from susceptible and resistant *B. glabrata* strains, showing that striking differences in proteins expressed, with susceptible snail haemocytes exhibiting extensive downregulation of protein expression and a lower level of constitutively expressed proteins involved immunity (e.g., *Bg*FREP2) and oxidation-reduction compared to resistant snails (Dinguirard et al., 2018). This could explain why BS-90 haemocytes alone can kill *S. mansoni* sporocysts. After challenge, the transcript abundance of numerous *Bg*FREPs increases by 3-fold or greater in BS-90 when compared to M-line *B. glabrata* (Adema et al., 1997). Noteworthy in the context of our current study is that following challenge BS-90 snails with *S. mansoni*, transcript abundance for *Bg*FREP2 increases over 50-fold by 1 day post-exposure (Dinguirard et al., 2018; Hertel et al., 2005), *Bg*FREP3 increases around 4-fold by 1 day post-exposure (Hanington et al., 2010), and *Bg*TEP1 which increases by 2- to 3.5-fold from 6 to 24 h post-exposure (Portet et al., 2018). These dramatic increases in the abundance of immune-relevant transcripts occurring over the first 12-24 h post exposure to *S. mansoni* sporocysts in BS-90 snails aligns closely with our results that demonstrate sporocyst killing by BS-90 haemocytes rapidly increases after 12 h post-exposure (Fig. 4).

Numerous studies have demonstrated the production of ROS by haemocytes in *B. glabrata*, especially H2O2, which plays a vital role in anti-schistosome defense (Galinier et al., 2013; Hahn et al., 2000, 2001). Haemocytes from *S. mansoni*-resistant snails produce significantly more ROS than susceptible snails (Bender, Broderick, Goodall, & Bayne, 2005; Bender, Goodall, Blouin, & Bayne, 2007; Goodall, Bender, Broderick, & Bayne, 2004). In our study, ROS was found to be critical in BS-90 haemocyte-mediated *S. mansoni* sporocyst killing, as this ability was significantly reduced after pre-incubation with the ROS inhibitors NAC and DPI (Fig. 4E). However, according to the phenomenon of passively transferred resistance, the cytotoxic potential of susceptible haemocytes may not fundamentally different from resistant snails (Granath & Yoshino, 1984). We infer that M-line plasma may lack some factors or the levels of these factors do not reach a certain threshold and fail to stimulate haemocytes to produce ROS, or M-line haemocytes are short of some receptors to receive signals that up-regulate ROS. Previously, our team identified a toll-like receptor (*Bg*TLR) from haemocyte surface of *B. glabrata* snails, which differed in abundance between M-line and BS-90 strains (Pila, Tarrabain, et al., 2016). It has been repeatedly demonstrated in other organisms that activation of Toll-like receptors is associated with increased intracellular ROS (Asehnoune, Strassheim, Mitra, Kim, & Abraham, 2004; Tsung et al., 2007; Wong et al., 2009).

Our results indicate that the *Bg*FREP3-*Bg*TEP1 complex plays a role in raising ROS levels in M-line haemocytes (Fig. 4F). We found that BS-90 haemolymph contains more *Bg*FREP3 protein than M-line (Fig. 1B). Moreover, the BS-90 haemocytes significantly increased the expression level of *Bg*FREP2 during encapsulating *S. mansoni* sporocysts compared with susceptible snail haemocytes (Dinguirard et al., 2018). These suggest that *Bg*FREP3 and *Bg*FREP2 may play a crucial role in inducing haemocytes which take part in encapsulating sporocysts to generate ROS. We suppose that *Bg*TEP1 also plays an opsonin role in the *B. glabrata* snail haemolymph, which interacts with molecules on the surface of haemocytes, thereby promoting the phagocytosis of pathogens.

Finally, our data hints at fundamental differences in the *Bg*FREP3-mediated immune response between BS-90 and M-line *B. glabrata*. Protein Pull-down experiments yielded two obvious and unique protein bands that were present in BS-90 but not M-line pull-downs. We identified the bands to be a variant of *Bg*FREP3.3 and *Bg*FREP2 (Fig. 1C), showing different immune factor interactomes between different strains. This suggests that *Bg*FREP3 variants and *Bg*FREP2 in BS-90 plasma are more prone to r*Bg*FREP3 binding, and the combination of different versions of *Bg*FREPs seems to play an important and unknown role in snail resistance. Both *Bg*FREP2 and *Bg*FREP3 are characterized by high diversity, our results imply that the versions of *Bg*FREP2 and *Bg*FREP3 in the two snail strains are different. After exposure to *S. mansoni*, the expression of *Bg*FREP2 in resistant snails is superior to that of susceptible snails (Dinguirard et al., 2018; Gordy et al., 2015; Hertel et al., 2005). This again suggests that *Bg*FREP2 is important in determining the outcome of the compatibility of *B. glabrata* against *S. mansoni*. Our study implies that *Bg*FREP3 seems to play an “all or none” switching effect in determining B.glabrata-*S. mansoni* compatibility, while *Bg*FREP2 seems to play an additive role.

It has been decades since the discovery and initial characterization of *Bg*FREPs in the *B. glabrata* immune response to *S. mansoni*. While numerous studies since have implicated both *Bg*FREPs (*Bg*FREP3 and *Bg*FREP2) along with a suite of other immune factors, as being important elements of the snail immune response to *S. mansoni*, how the response is coordinated has remained elusive. Our study presents a model in which many of the known determinants of snail resistance to *S. mansoni* are found to work in concert to protect the snail host against infection. We map a mechanism for the interaction of *Bg*FREP3, *Bg*TEP1, *Bg*FREP2 and Biomphalysin with *S. mansoni* sporocysts in *B. glabrata* snail haemolymph that ultimately leads to sporocyst killing (Fig. 5). Central to this response is *Bg*FREP3, which is able to recruit unique representatives of both *Bg*FREP2 and *Bg*FREP3 in *S. mansoni*-resistant BS-90 *B. glabrata*. With this new-found understanding of the *B. glabrata* immune response comes the ability to now undertake comprehensive assessments of how modifications to both the snail host and the parasite influence the underpinning immunological drivers of compatibility in this important model for studying human schistosomes.

**Figure 5.**
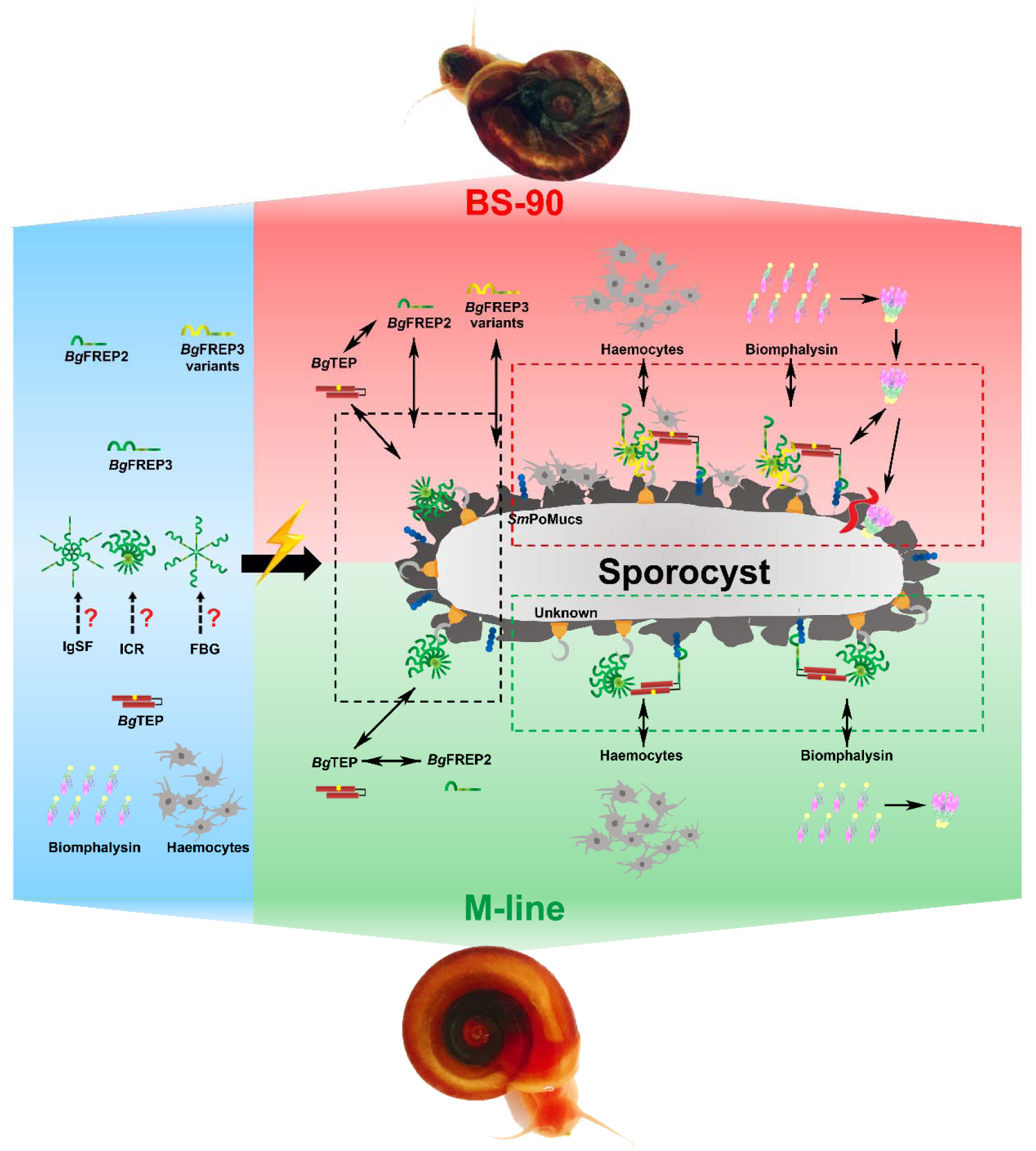
Interaction of *Bg*FREP3, *Bg*FREP2, *Bg*TEP1 and Biomphalysin in *B. glabrata* snail hemolymph and possible functional mechanisms against *S. mansoni* infection. This figure is a schematic drawing based on the results of previous studies and our work.

## Materials and Methods

### Live material and Experimental Treatments

M-line and BS-90 strain *B. glabrata* snails were used in this study. The M-line strain is highly susceptible to the PR-1/ NMRI strain *S. mansoni* infection (Newton, 1955), while the BS-90 strain is resistant. M-line and BS-90 strain *B. glabrata* snails, and *S. mansoni* were maintained at the University of Alberta as described previously (Pila, Gordy, et al., 2016). NMRI strain of *S. mansoni* was obtained from infected Swiss-Webster mice provided by the NIH/NIAID Schistosomiasis Resource Center at the Biomedical Research Institute (Cody et al., 2016). All animal work observed ethical requirements and was approved by the Canadian Council of Animal Care and Use Committee (Biosciences) for the University of Alberta (AUP00000057).

### Recombinant *Bg*FREP3, *Bg*TEP1 and *Bg*FREP2 Synthesis and Purification

Recombinant *Bg*FREP3 (r*Bg*FREP3, derived from GenBank: AY028461.1), *Bg*TEP1 (r*Bg*TEP1, GenBank: HM003907.1) and *Bg*FREP2 (r*Bg*FREP2, GenBank: AY012700.1) were generated by using the Gateway cloning system according to the manufacturer’s instructions (Life Technologies) as previously described (Hambrook, Kabore, Pila, & Hanington, 2018; Pila, Gordy, et al., 2016; Pila, Peck, & Hanington, 2017; Pila, Tarrabain, et al., 2016), with primer information as shown in Table. S1 and Sf9 cell codon-optimized *Bg*FREP3 Genscript as shown in Fig. S1. Briefly, the coding regions were amplified with Phusion high-fidelity DNA polymerase from targeted DNA templates in pUC57 plasmids (synthesized by GenScript). Blunt-end PCR products were cloned into the pENTR/D-TOPO vectors to generate entry clones. Plasmids from these entry clones were extracted and cloned into pIB/V5-His-DEST vectors in a Clonase recombination reaction to produce the expression clones. Recombinant plasmids were extracted from the expression clones and then transfected into Sf9 insect cells using Cellfectin reagent (Life Technologies). Cells that survived the screening of the antibiotic blasticidin (Thermo Fisher Scientific) were collected and lysed. Protein expression of r*Bg*FREP3, r*Bg*TEP1 and r*Bg*FREP2 were detected by Western blot using monoclonal antibody against the V5 tag (Thermo Fisher Scientific) on the recombinant proteins.

Sf9 cells stably expressing recombinant r*Bg*FREP3, r*Bg*TEP1 and r*Bg*FREP2 were selected, some of them were frozen as a seed stock and the rest were maintained in a medium containing blasticidin. Cultures of Sf9 cells expressing recombinant proteins were scaled up, and the pellet (containing r*Bg*FREP3) or the medium (containing secreted r*Bg*TEP1 and r*Bg*FREP2) was collected. Recombinant proteins were purified from the cell pellet lysate or the culture medium with HisTrap FF (GE Healthcare) columns using fast protein liquid chromatography (ÄKTA Pure, GE Healthcare). Purified r*Bg*FREP3, r*Bg*TEP1 and r*Bg*FREP2 were dialyzed against PBS buffer twice for 2 h each and then once overnight using Slide-A-Lyzer dialysis kit (Thermo Scientific).

### Preparation of *B. glabrata* plasma and pull-down experiment

Hemolymph was obtained from M-line and BS-90 strain *B. glabrata* snails by the head-foot retraction method (Pila, Gordy, et al., 2016). Upon collection, hemolymph from ∼5 snails of each strain (8–12 mm in diameter) was immediately placed in 1.5 mL tubes on ice, EDTA was added to a final concentration of 150 μg/mL. Samples were centrifuged twice at 20,000 × g for 15 min and the cell-free fraction was collected. The 4 M Imidazole Stock Solution was added to a final concentration of 10 mM imidazole. Plasma samples were used immediately for subsequent pull-down experiment.

Pull-down assays were conducted by using Pull-Down PolyHis Protein:Protein Interaction Kit (Thermo Scientific) according to the manufacturer’s instructions with some modifications. Briefly, first we prepared the “bait” proteins, r*Bg*FREP3 and r*Bg*TEP1, from previously purified proteins or Sf9 cell lysate expressing r*Bg*FREP3 or concentrated medium containing r*Bg*TEP1. We then loaded the HisPur Cobalt Resin equilibrated for at least 30 minutes at room temperature onto the column according to the kit instructions. The prepared polyhistidine-tagged r*Bg*FREP3 and r*Bg*TEP1 was immobilized on the columns after five times washing with wash solution [1:1 of Tris-buffered saline (TBS) solution:Pierce Lysis Buffer containing 10 mM imidazole]. For cell lysates or concentrated medium containing r*Bg*FREP3 and r*Bg*TEP1, protein expression was previously analyzed by SDS-PAGE and Western blot, and 800 μL was added; for purified r*Bg*FREP3, at least 150 µg (about 500 μL) was added. After incubation for 2 h at room temperature or overnight at 4°C, the columns were washed 5-8 times with wash solution and then 800 µL of prepared plasma (prey) containing 10 mM imidazole was added to the columns. The columns were incubated at 4°C for overnight with gentle rocking motion every few h. After that, the columns were washed 5-8 times with the wash solution, and then eluent was added and incubated at room temperature for 30 min with gentle rocking on a rotating platform.

The eluent collected by centrifugation was used for SDS-PAGE analysis and silver staining (Pierce^™^ Silver Stain for Mass Spectrometry, Thermo Scientific) was performed according to the manufacturer’s instructions. After silver staining, we excised the desired bands for LC-MS/MS to identify the proteins.

### Mass spectrometry analysis

Mass spectrometry and protein identification was completed by Alberta Proteomics and Mass Spectrometry Facility at University of Alberta. In-gel trypsin digestion was performed on the samples. Briefly, the excised gel bands were reduced (10 mM mercaptoethanol in 100 mM bicarbonate) and alkylated (55 mM iodoacetamide in 100 mM bicarbonate). After dehydration enough trypsin (6 ng/μL, Promega Sequencing grade) was added to just cover the gel pieces and the digestion was allowed to proceed overnight (∼16 h) at room temperature. Tryptic peptides were first extracted from the gel using 97% water/2% acetonitrile/1% formic acid followed by a second extraction using 50% of the first extraction buffer and 50% acetonitrile.

The digested samples containing tryptic peptides were resolved and ionized by using nanoflow HPLC (Easy-nLC II, Thermo Scientific) coupled to an LTQ Orbitrap XL hybrid mass spectrometer (Thermo Scientific). Nanoflow chromatography and electrospray ionization were accomplished by using a PicoFrit fused silica capillary column (ProteoPepII, C18) with 100 μm inner diameter (300 Å, 5 μm, New Objective). The mass spectrometer was operated in data-dependent acquisition mode, recording high-accuracy and high-resolution survey Orbitrap spectra using external mass calibration, with a resolution of 30,000 and m/z range of 400–2,000. The fourteen most intense multiply charged ions were sequentially fragmented by using collision induced dissociation, and spectra of their fragments were recorded in the linear ion trap; after two fragmentations all precursors selected for dissociation were dynamically excluded for 60 s. Data was processed using Proteome Discoverer 1.4 (Thermo Scientific) and a *B. glabrata* proteome database (UniProt) was searched using SEQUEST (Thermo Scientific). Search parameters included a precursor mass tolerance of 10 ppm and a fragment mass tolerance of 0.8-Da. Peptides were searched with carbamidomethyl cysteine as a static modification and oxidized methionine and deamidated glutamine and asparagine as dynamic modifications.

### Sporocyst transformations of *S. mansoni* and immunofluorescence staining

Miracidia were obtained from eggs isolated from the livers of mice infected with *S. mansoni* (NMRI) as described previously (Hambrook et al., 2018). Briefly, newly hatched miracidia were collected for cultivation in Chernin’s Balanced Salt Solution (CBSS) (Chernin, 1963) containing 1 g/L each of glucose and trehalose as well as 1% pen-strep antibiotics (Life Technologies). After cultivation for 24 h at 27°C under normoxic conditions, most miracidia transformed to primary sporocysts. Primary sporocysts were washed five times with snail PBS (sPBS) (Yoshino & Laursen, 1995) and transferred to 15 mL centrifuge tubes (Corning). All in-tube washes were performed at 4°C and sporocysts were pelleted by centrifugation for 2 min at 300 × g.

After overnight 2% paraformaldehyde/sPBS (pH 7.2) fixation, sporocysts were washed five times with sPBS followed by 1 h incubation in 5% BSA/0.02% azide/sPBS (blocking buffer) at room temperature. Sporocysts was grouped and incubated with r*Bg*FREP3 (500 ng/μL), r*Bg*TEP1 (50 ng/μL) and r*Bg*FREP2 (150 ng/μL) in blocking buffer overnight at 4°C; control group were incubated in blocking buffer without recombinant proteins. Following five washes with 1% BSA/sPBS, sporocysts were treated with anti-V5 antibody (Life Technologies) diluted 1:400 in blocking buffer. Following primary antibody treatment, sporocysts were washed five times with 1% BSA/sPBS and incubated with a solution containing 7.5 U/mL Alexa Fluor 488 phalloidin (Invitrogen) and 4 µg/mL Alexa Fluor 633 goat anti-mouse secondary antibody (Invitrogen) in blocking buffer overnight at 4°C. Finally, sporocysts were washed five times with 1% BSA/sPBS, dropped into a drop of DAPI staining solution, incubated for 5 min at room temperature, and transferred onto microscope slides. Preparations were then covered with a cover glass and sealed with nail polish to prevent desiccation. Slides were imaged using an Axio imager A2 microscope (Zeiss), and analyzed using Zen 2011 software (Zeiss) and Photoshop CS4 (Adobe Systems Inc., USA).

### Anti-*Bg*FREP3 polyclonal antibody generation and validation

The IgSF1 and FBG domains of *Bg*FREP3 were screened for regions of presumed antigenicity using the OptimumAntigen^TM^ design tool (GenScript). Three polyclonal antibodies were generated specific for different peptides (14-15 amino acids long, ∼ 4.4-kDa) targeting each domain. The information of these peptides is: anti-IgSF #1: CSFKKDDLSDSKQRS; anti-IgSF #2: CHNKYSEGRIDKSSN; anti-IgSF #3: CNINKDLDFKEQNIT; anti-FBG #1: TTFDRDNDEYSYNC; anti-FBG #2: GWKEYRDGFGDYNIC and anti-FBG #3: CLNGKWGSSDFAKGV. Peptides were synthesized by GenScript and used to immunize rabbits. Rabbits received a primary immunization and two secondary boosts at 2 and 5 weeks. One week after the second boost, rabbits were exsanguinated and the serum IgG was affinity-purified using protein A/G affinity column. These were further purified using affinity resin conjugated with the immunization peptides. The anti-*Bg*FREP3 antibodies used in this study were effective for Western blot detections at a concentration of 1:1,000.

The purified antibodies were supplied by GenScript along with synthesized peptides and pre-immune serum. Specificity of the antibodies was tested against the respective peptides as well as snail plasma and haemocyte lysates. Dot blots were initially used to determine whether the antibodies recognized their peptides as well as the specificity of the recognition. Briefly, 2 µL of each supplied peptide was slowly pipetted in one spot on a nitrocellulose membrane that was divided into a grid with a pencil. The deposited peptides were dried at room temperature for about 10 min, creating a spot of approximately 2–3 mm. The membrane was then incubated in blocking buffer and processed using the same procedure described for Western blot following sample transfer to nitrocellulose membranes. In order to determine that the peptides were not being recognized by any pre-immune serum components, the above procedure was followed exactly with the exception that the respective pre-immune sera were used for the primary incubations in place of antibodies.

### Western and far-Western blot analysis

To test whether r*Bg*TEP1 complexes with r*Bg*FREP3 and r*Bg*FREP2, purified r*Bg*TEP1 was incubated with r*Bg*FREP3 or r*Bg*FREP2 in the following ratios (wt/wt in µg): 1:1, 1:0.75, 1:0.5, and 1:0.25 in 1.5 mL tubes for 2 h at room temperature on a Labquake shaker (ThermoFisher Scientific) to allow potential complexes to form. After the incubation, samples were subjected to western blot detection under denaturing (Laemmli protein loading buffer with β-mercaptoethanol and heated at 95°C for 10 min) and non-denaturing (no β-mercaptoethanol nor heating) conditions, respectively.

The Western blot experiment process was as described previously (Pila, Peck, et al., 2017). Samples were loaded on 6-14% (vol/vol) SDS/PAGE gels and run on the Mini PROTEAN Tetra system (Bio-Rad) at 150 V for 1.5-2 h. Samples were then transferred for 1.5 h onto 0.45 μm supported nitrocellulose membranes (Bio-Rad). Blocking was done for 1 h at room temperature in 5% (wt/vol) skimmed milk or bovine serum albumin (BSA) prepared in TBS solution plus 0.1% Tween-20 (TBS-T buffer) before staining for 1 h in anti-V5 mouse primary antibody at a dilution of 1:5,000 in blocking buffer. Membranes were washed in TBS-T buffer for 10 min, then twice for 5 min each and once in TBS solution for 5 min. Membranes were then incubated for 1 h in HRP-conjugated rabbit anti-mouse antibody diluted 1:5,000 in blocking buffer followed by a wash step as described above. Finally, the blot was developed by incubating the membranes in SuperSignal West Dura Extended Duration substrate (Thermo Scientific). Chemiluminescent signals were acquired on the ImageQuant LAS 4000 machine (GE Healthcare).

To detect the *Bg*FREP3 background expression in the *B. glabrata* snail hemolymph, the hemolymph of four snails was obtained from each of M-line and BS-90 strain *B. glabrata* snails. The hemolymph from each snail was diluted twice with lysis buffer (50 mM Tris, pH 7.8; 150 mM NaCl; 1% Nonidet P-40 and 150 μg/mL EDTA). After incubation for 30 minutes on ice, samples were centrifuged at 20,000 × g for 15 min and the supernatant was collected. After quantification, samples (160 μg/well) were used for subsequent Western blot analysis. The primary antibodies were polyclonal antibodies raised in rabbits against the IgSF and FBG domains of r*Bg*FREP3 (1:500 dilution) and the secondary antibody was a HRP-conjugated goat anti-rabbit IgG (1:5000 dilution).

Far-Western blotting is used to probe a membrane containing transferred protein with another protein to detect specific protein-protein interactions (Y. Wu, Li, & Chen, 2007). In our study, we detected r*Bg*TEP1 (“prey” protein) separated by SDS-PAGE with the biotinylated r*Bg*FREP3 and r*Bg*FREP2 (“bait” or “probe” proteins) and a streptavidin-HRP chemiluminescent detection system. If r*Bg*TEP1 forms a complex with r*Bg*FREP3 or r*Bg*FREP2, r*Bg*FREP3 would be detected on spots in the membrane where r*Bg*TEP1 was located. First, r*Bg*FREP3 and r*Bg*FREP2 was biotinylated with EZ-Link Sulfo-NHS-LC-Biotin (Thermo Scientific) according to the manufacturer’s instructions. Recombinant *Bg*TEP1 was separated by SDS-PAGE, and transferred to a membrane, as in a standard western blot. Recombinant *Bg*TEP1 in the membrane was then denatured and renatured as described previously (Y. Wu et al., 2007). The membrane was then blocked and probed with biotinylated r*Bg*FREP3 or r*Bg*FREP2. After washing as described above in a standard western blot, the membrane was incubated with streptavidin-HRP diluted 1:100,000 in blocking buffer at room temperature for 1 h. Chemiluminescent signals were detected using the ImageQuant LAS 4000 machine (GE Healthcare).

### Quantitative real-time PCR analysis of *Bg*TEP1 expression

A TRIzol reagent was utilized to extract total RNA from 5 whole unchallenged snails from each strain independently, after which RNA was purified using a PureLink RNA mini kit (Life Technologies). A NanoVue spectrophotometer (ThermoFisher Scientific) was used to determine RNA concentration, and first-strand cDNA synthesis using the qScript cDNA synthesis kit (Quanta Biosciences) was performed using 1 μg of RNA template. The cDNA was diluted fivefold, and 5 µL was used as template in RT-PCR using primers (Table S1) specific for *Bg*TEP1 and *B. glabrata* β-actin (*Bg*Actin) in a SYBR Green detection system (PerfeCTa SYBR Green FastMix; Quanta Biosciences). All RT-PCRs were performed on a QuantStudio 3 PCR system (Applied Biosystems) using the following thermocycling conditions: initial hold at 95°C for 10 min, followed by 40 cycles of 95°C for 15 s and 60°C for 1 min, with data collection every cycle. Specificity for the gene specific primers RT-PCR was confirmed by continuous melt curve analysis.

### *In vitro S. mansoni* sporocyst killing assays

Haemolymph from 25 M-line or BS-90 strain *B. glabrata,* sterilized as reported in Hahn *et al*., 2001 (Hahn et al., 2001), was pooled into 1.5 mL tubes and kept on ice. Haemocytes were isolated from plasma by centrifugation at 70 × g for 10 min at 4°C followed by aspiration of the cell free plasma, which was kept for assessment of plasma-mediated killing of sporocysts. Haemocytes were resuspended in modified *Bg*e-cell medium (m*Bg*e) medium [22% Schneider’s Drosophila medium (Thermo Scientific), 7 mM D-glucose, 24 mM NaCl, and 20 mg/mL gentamicin at pH 7.4] and washed three times following the same centrifugation protocol. Haemocytes were then resuspended to a final concentration of 50 cells/µL in m*Bg*e medium and 200 µL of this suspension was added to the wells of a 96-well tissue culture plate pre-treated with poly-L-lysine that contained 10 *S. mansoni* sporocysts that were transformed using previously published protocols (Hambrook et al., 2018). Treatment consisted of 200 pM of each recombinant protein (*Bg*FREP3, *Bg*TEP1, *Bg*FREP2). Three anti-*Bg*FREP3 antibodies (anti-IgSF #1 and #3, anti-FBG #1) that were generated to peptides within the IgSF and FBG domains were used as a cocktail (combined concentration of 2.5 mg/mL) were used the antibody blocking assays. Sporocyst viability was assessed using the protocol published by Hahn *et al*., 2001 (Hahn et al., 2001). Briefly, Propidium iodide (Millipore Sigma), was added to each well to a final concentration of 10 µg/mL. Sporocyst viability was assessed using a fluorescent inverted microscope (Zeiss) starting at 3 h post incubation with haemocytes/plasma and then measured again at 6, 12, 24, 36 and 48 h post incubation. Sporocyst death was determined by propidium iodide-staining of the nuclei of the sporocyst. Any sporocyst deemed dead, was left in the well until the end of the time-course so that the experimental conditions were not disturbed differentially in wells where sporocyst mortality was more frequent.

The pooled plasma from M-line and BS-90 snails. Cell-free plasma was ultracentrifuged at 30,000 rpm for 3 h at 4°C to remove haemoglobin, which is known to have a toxic effect on sporocysts (Bender et al., 2002). Ultracentrifuged plasma was mixed 50:50 with m*Bg*e medium at room temperature and then incubated with 10 *S. mansoni* sporocysts according to the protocol outlined above.

### Assessment of reactive oxygen species production by haemocytes *in vitro*

To assess the role that ROS had on sporocyst killing by haemocytes the above haemocyte *in vitro* killing assays were repeated with addition of two well characterized inhibitors of ROS; diphenyleneiodonium chloride (DPI) and N-acetyl-L-cysteine (NAC). Both ROS inhibitors (DPI at a final concentration of 150 nM and NAC at a final concentration of 1 mM) were incubated along with haemocytes and *S. mansoni* sporocysts as described above in the haemocyte-based sporocyst killing assay.

### Bioinformatic and Statistical analysis

We used Clustal Omega (https://www.ebi.ac.uk/Tools/msa/clustalo/) to align the amino acid sequences for *Bg*TEPs, Biomphalysins and *Bg*FREPs which acquired from GenBank [National Center for Biotechnology Information (NCBI)]. The alignment files were downloaded and visually inspected for obvious inaccuracies, modified if need be, and then annotated based on information provided by NCBI. To show the distribution of the identified peptides over a full length sequence, the amino acid sequences of *Bg*TEP1.5 (ADE45333.1), Biomphalysin (A0A182YTN9) and *Bg*MFREP2 (AAK13550.1) were imported into Vector NTI Advance 11.5 (Thermo Fisher Scientific) and the identified peptides were annotated. Images produced by Vector NTI software were saved by FSCapture and further processed with FSCapture and Adobe Photoshop CS4. To determine significant differences in sporocysts killing assays, one-way ANOVA with Tukey’s post hoc tests were performed using GraphPad Prism version 6.0f for Mac OS X (GraphPad; www.graphpad.com). Statistical significance threshold was set at P ≤ 0.05.

## Acknowledgements

*B. glabrata* snails provided by the NIAID Schistosomiasis Resource Center of the Biomedical Research Institute (Rockville, MD) through NIH-NIAID Contract HHSN272201700014I for distribution through BEI Resources. These studies were supported by funds provided by the Natural Sciences and Engineering Research Council of Canada #2018-05209 and 2018-522661 (PCH), and National Natural Science Foundation of China (NSFC 31272682), Guangxi 16 Natural Science Foundation (Key Project, 2016JJD130059) Special Fund for Team Building, Beibu Gulf University (Former Qinzhou University, 2015).

## Author Contributions

H.L. and P.C.H. designed research; H.L., J.R.H., E.A.P., A.A.G., and J.F. performed research; H.L., J.R.H., E.A.P., X.W. and P.C.H. analyzed data; and H.L., J.R.H., E.A.P., A.A.G., and P.C.H. wrote the paper.

## Supplemental figure titles and legends

**Figure 1—figure supplement 1.**
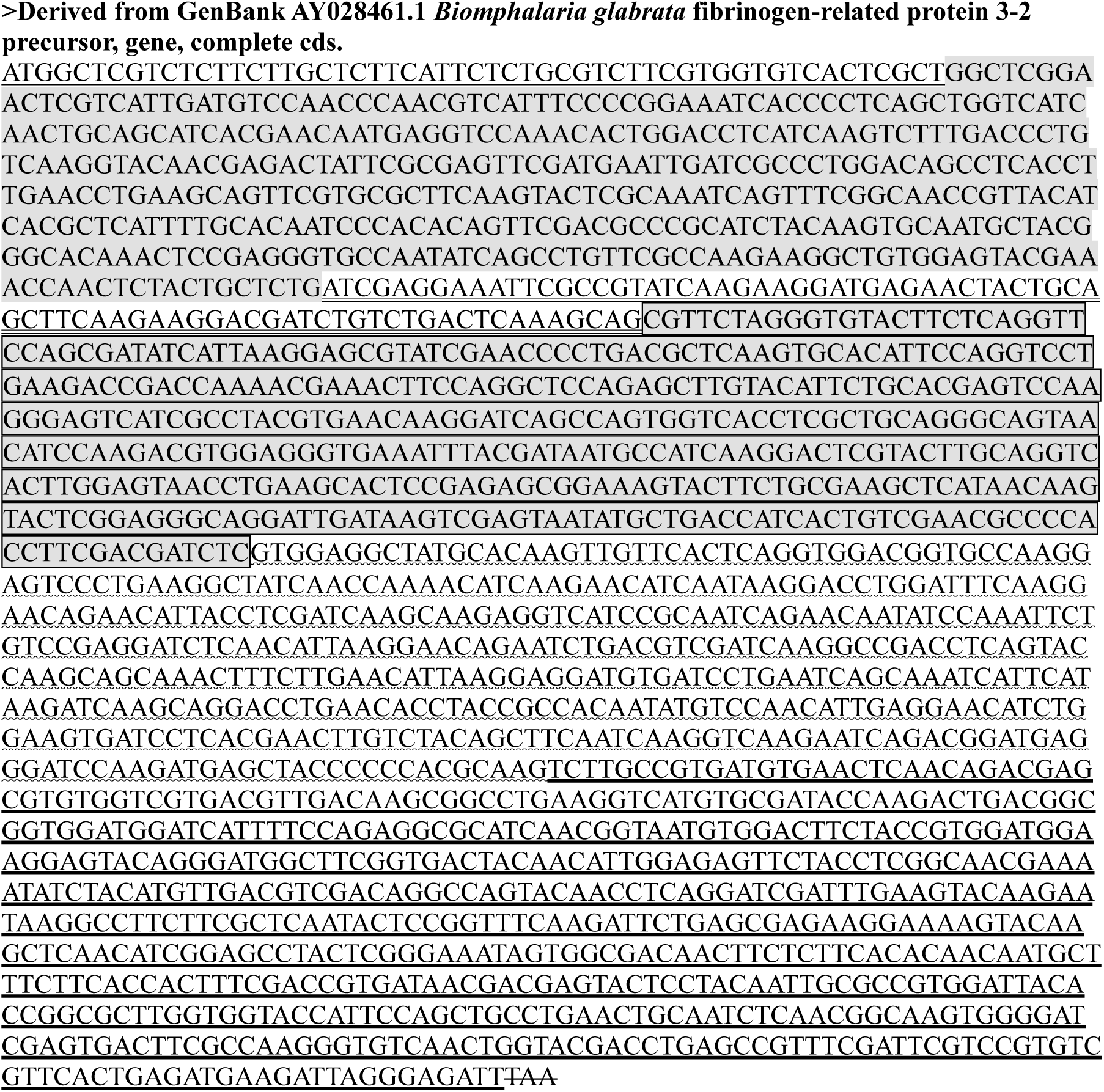
Nucleotide sequence for cloning r*Bg*FREP3. This sequence is derived from GenBank AY028461.1. There are differences in nucleotide sequence but the amino acids match 100%. The differences are due to codon-optimization for expression in insect Sf9 cells. GenBank annotations are used to locate the individual *Bg*FREP3.2 domains: underline, signal peptide; gray shading, IgSF1 domain; double underline, small connecting region; gray shading plus frame, IgSF2 domain; wavy underline, ICR; bold underline, FBG domain; delete line, termination codon.

**Figure 1—figure supplement 2.**
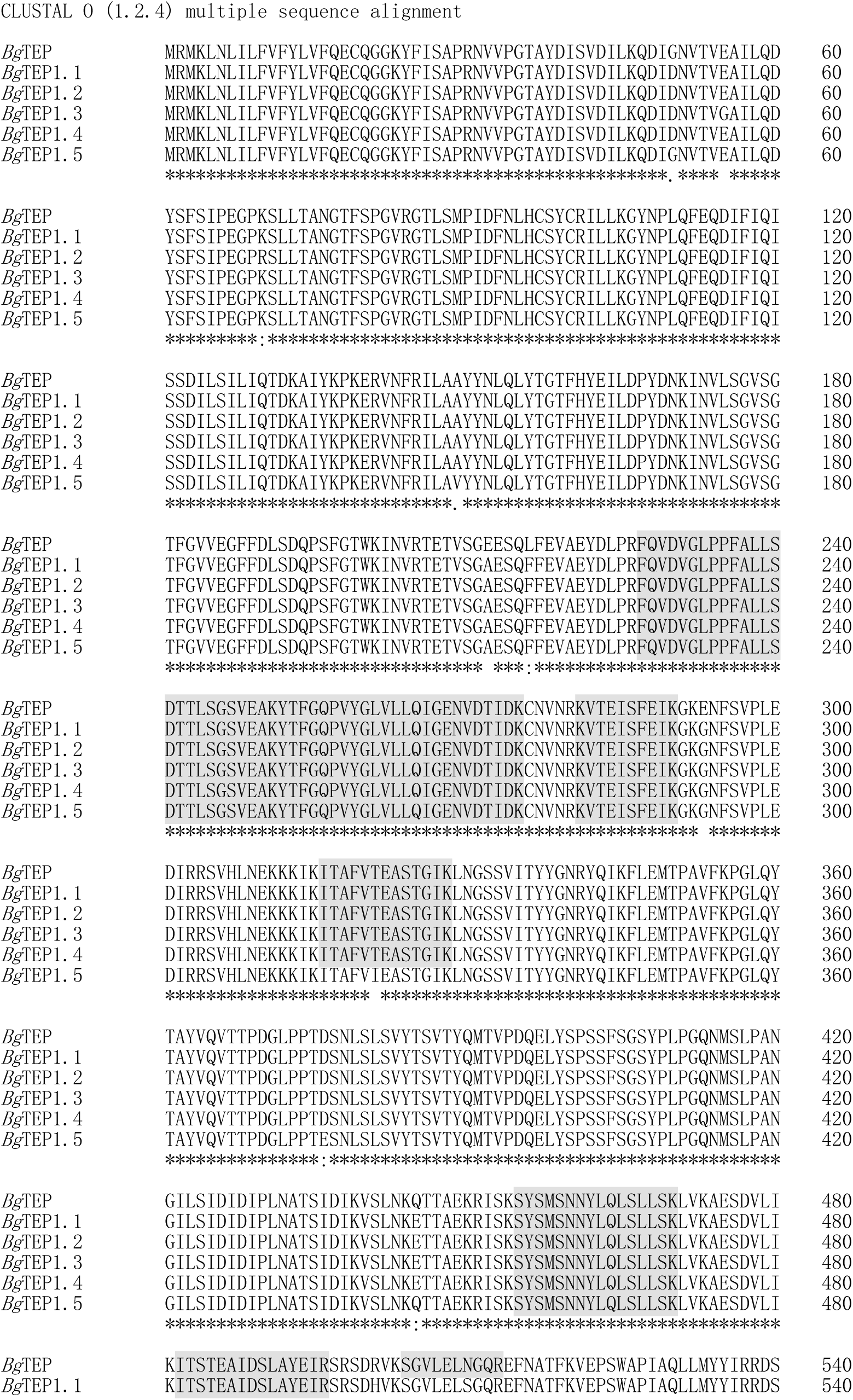
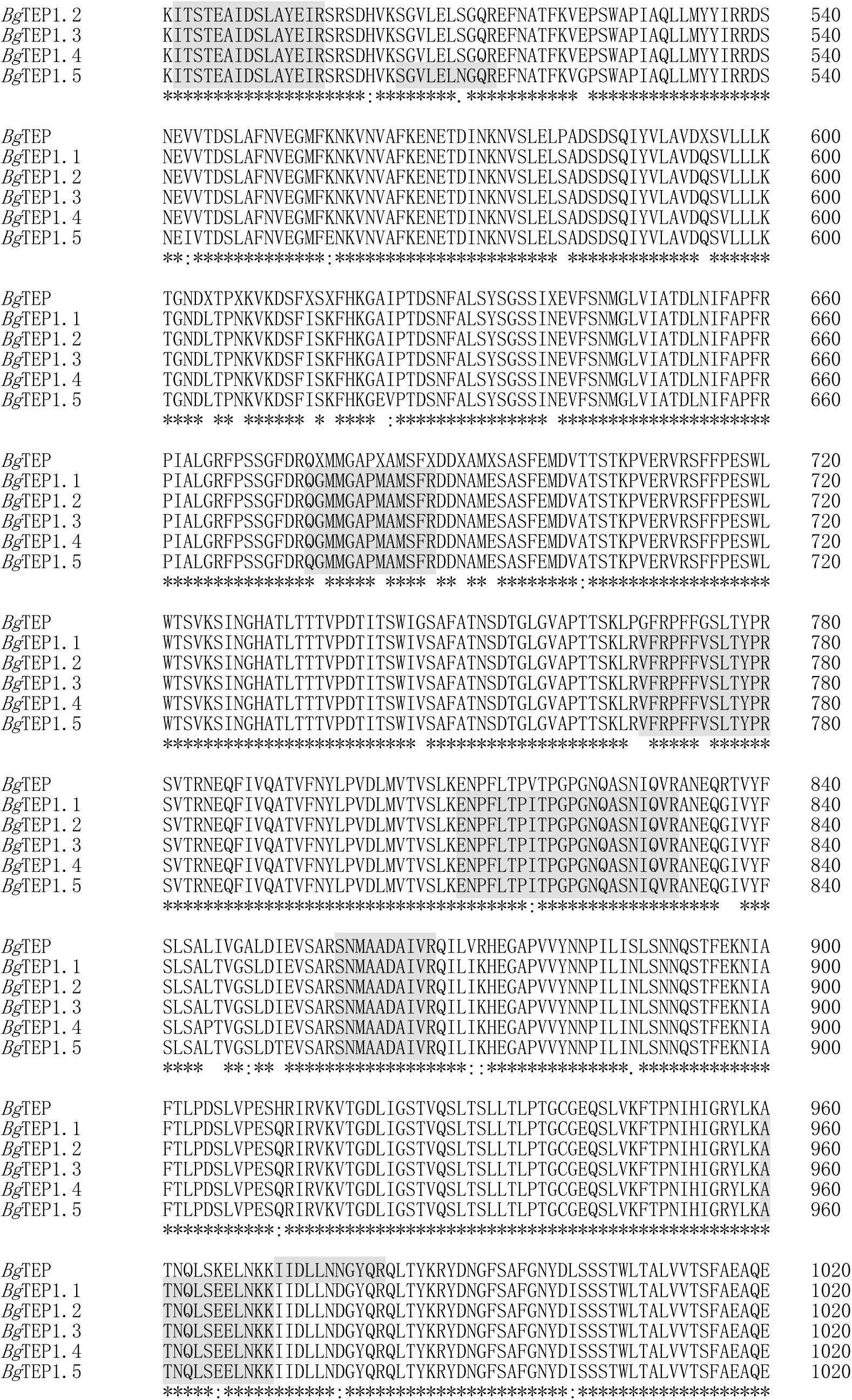
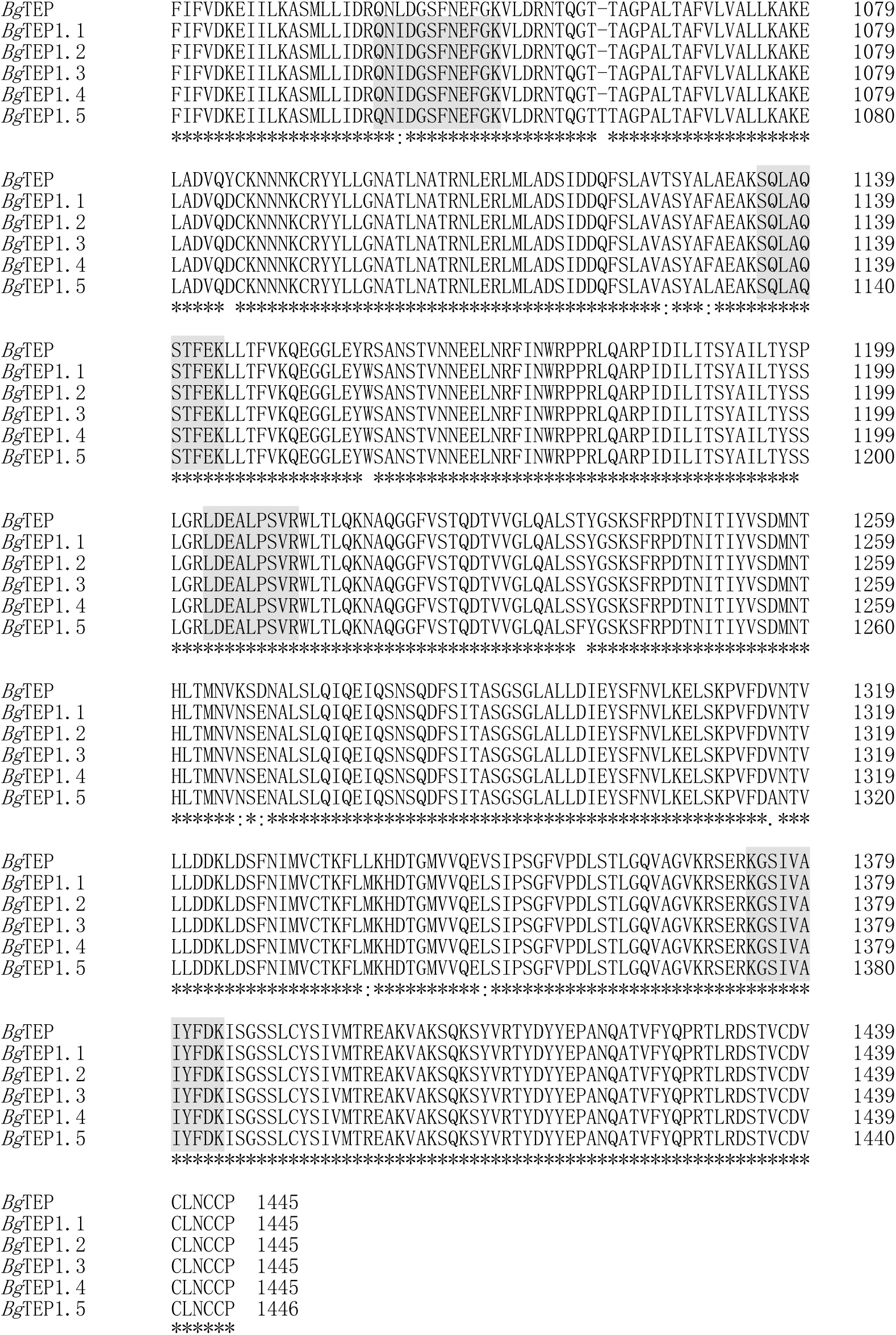
Alignment of multiple *Bg*TEP amino acid sequences and distribution of identified peptides. Peptides identified by LC-MS/MS from r*Bg*FREP3 pull-down experiments are highlighted in gray. The GenBank accession numbers of each entry are: *Bg*TEP, ACL00841.1; *Bg*TEP1.1, ADE45332.1; *Bg*TEP1.2, ADE45339.1; *Bg*TEP1.3, ADE45340.1; *Bg*TEP1.4, ADE45341.1 and *Bg*TEP1.5, ADE45333.1.

**Figure 1—figure supplement 3.**
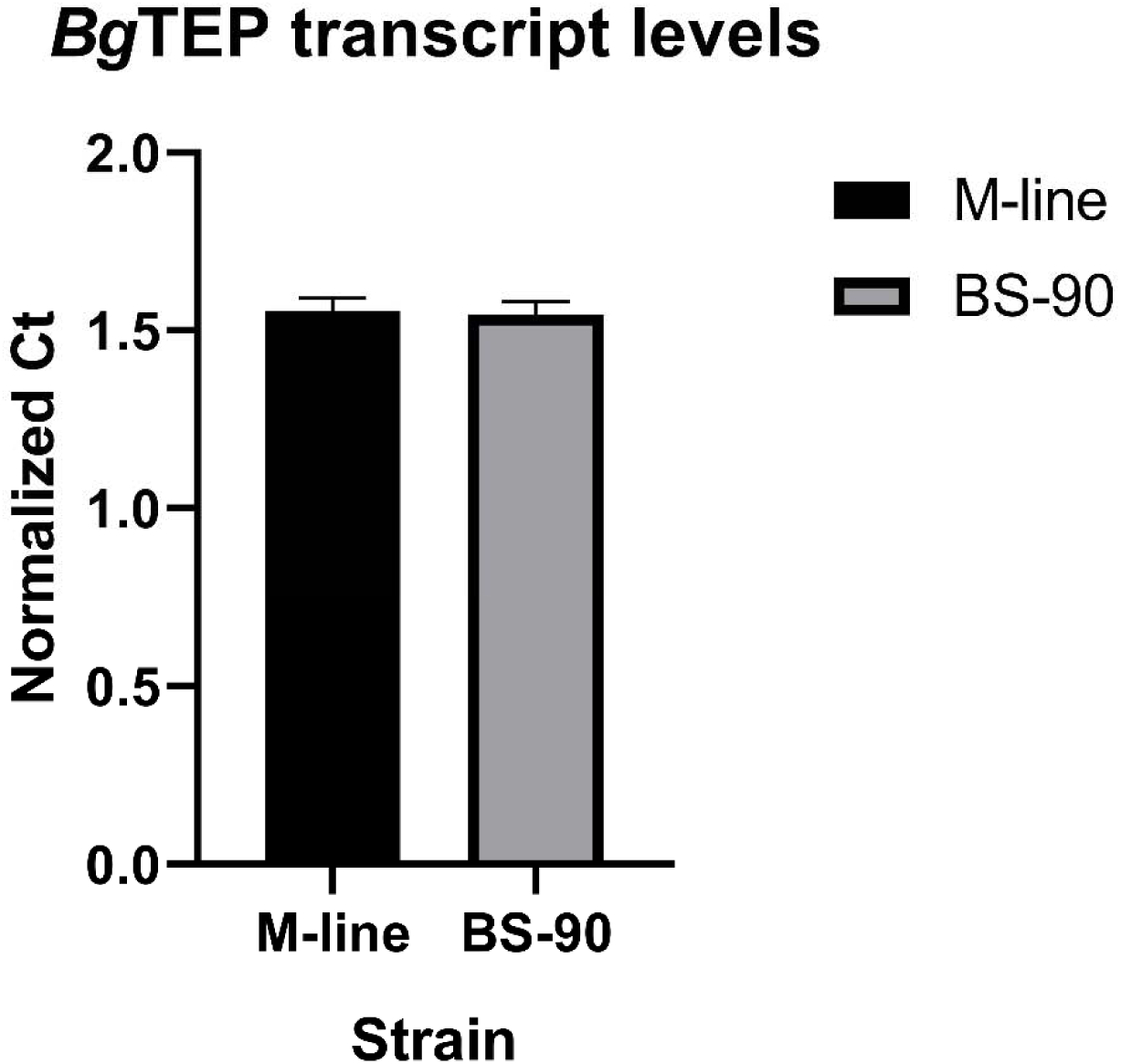
Normalized transcript abundance of *Bg*TEP1 in unchallenged M-line (black bar, n=5) and BS-90 (grey bar, n=5) snails indicating a lack of significant difference (p > 0.05) between the two strains.

**Figure 1—figure supplement 4.**
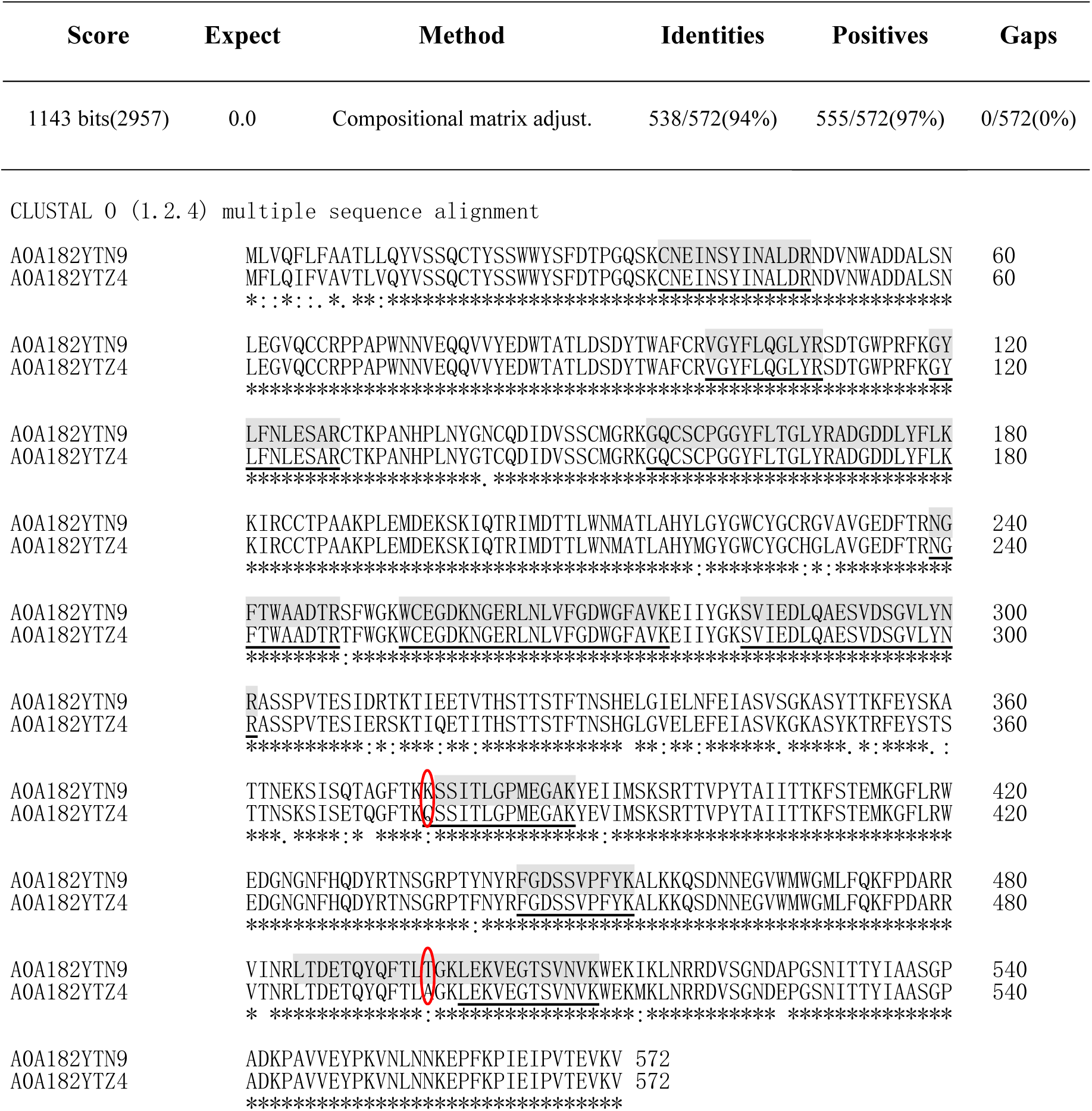
The alignments of two Biomphalysin variants (UniProtKB/TrEMBL: A0A182YTN9 and A0A182YTZ4) identified by LC-MS/MS. Peptides identified from A0A182YTN9 and A0A182YTZ4 are highlighted in gray and underline, respectively. Two red oval boxes representing differences in peptide sequences that distinguish two Biomphalysin variants.

**Figure 1—figure supplement 5.**
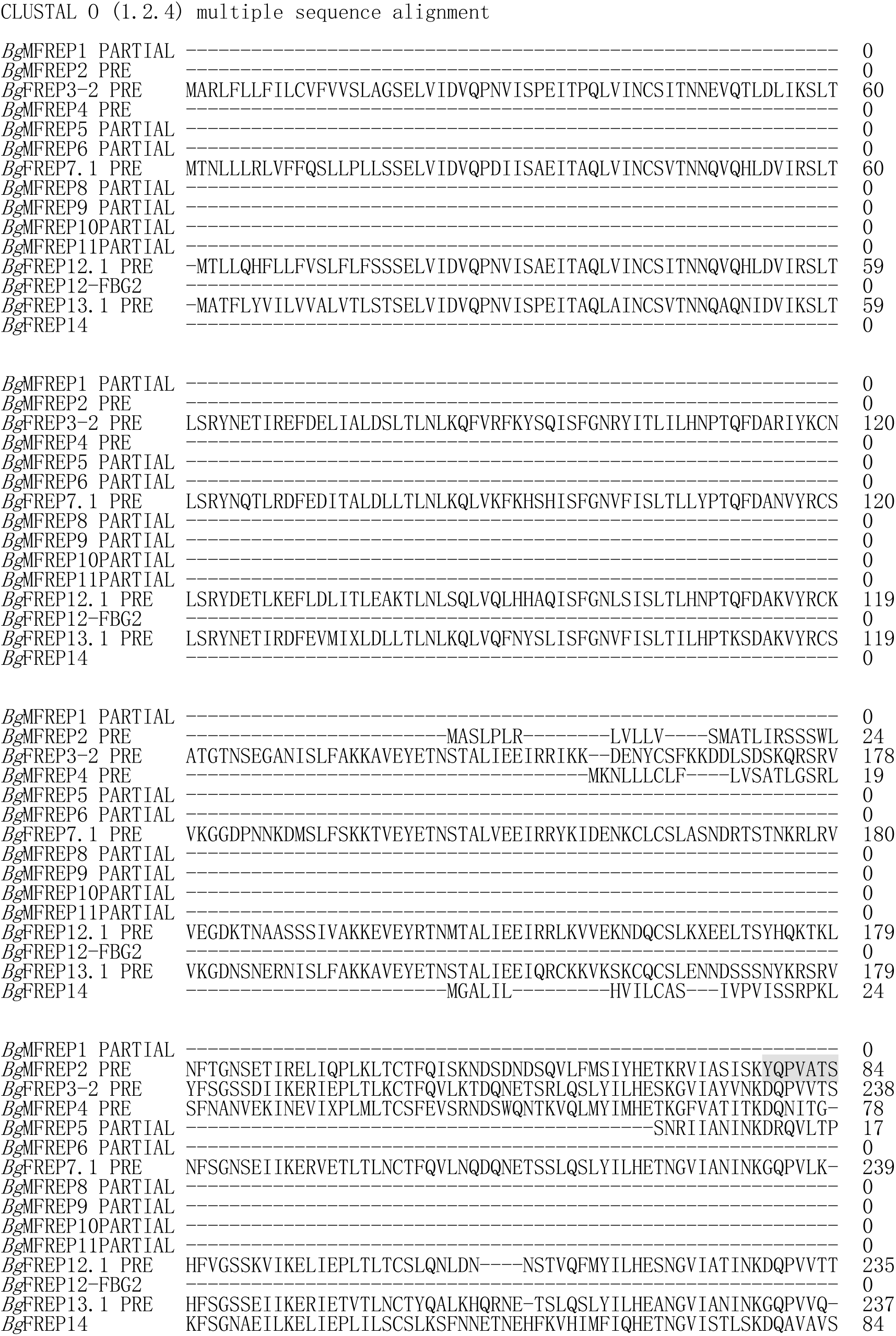
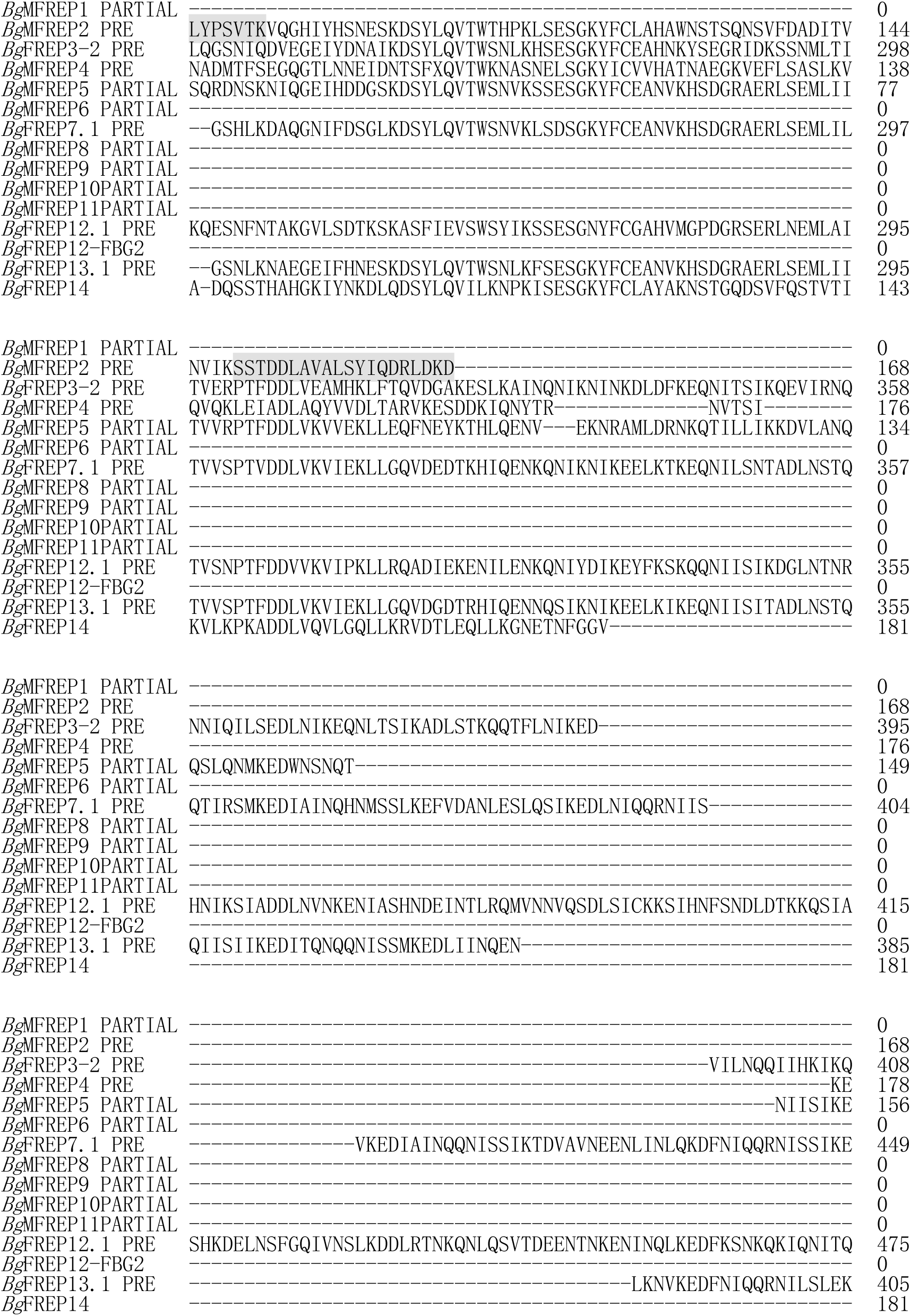
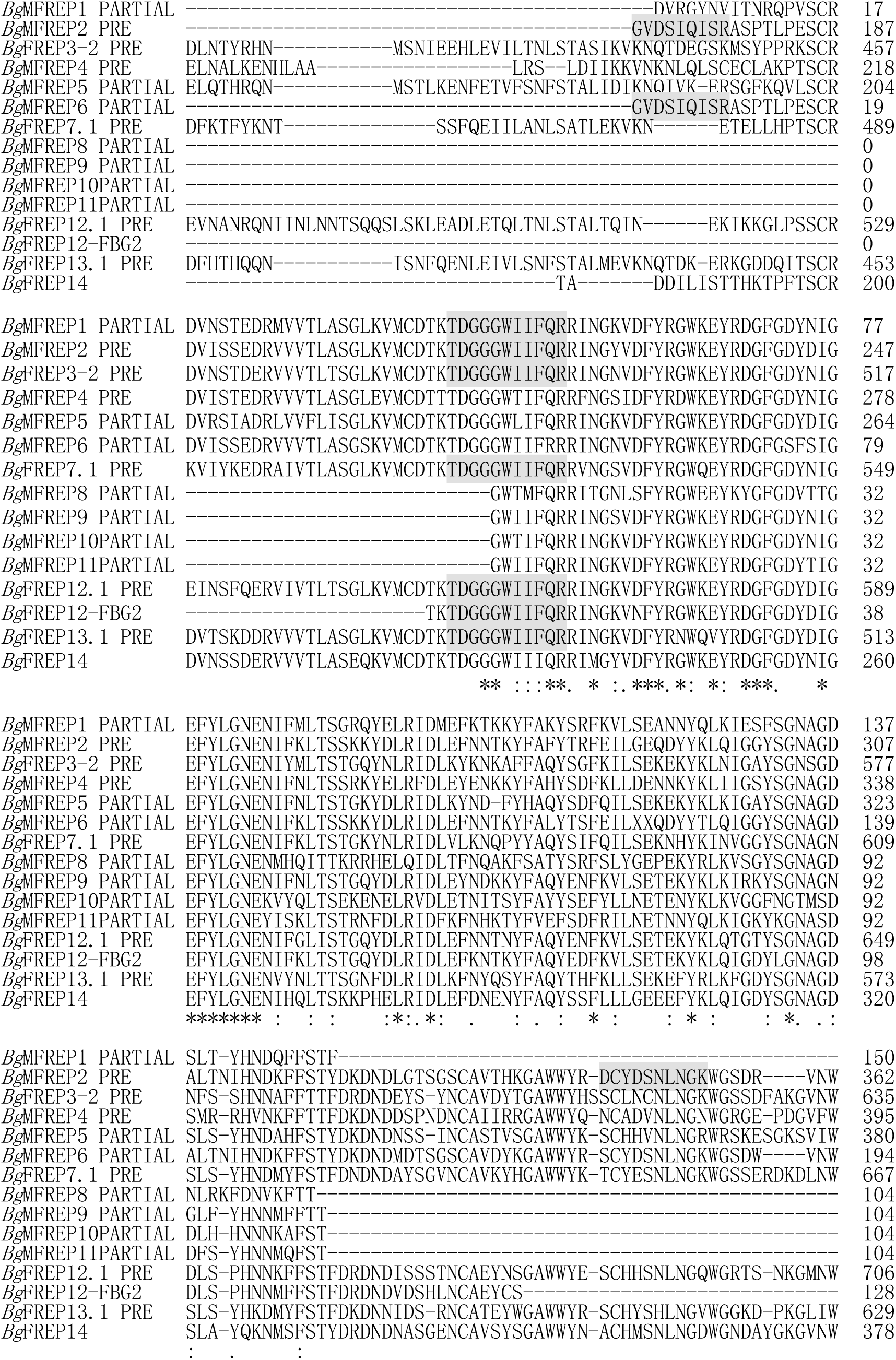
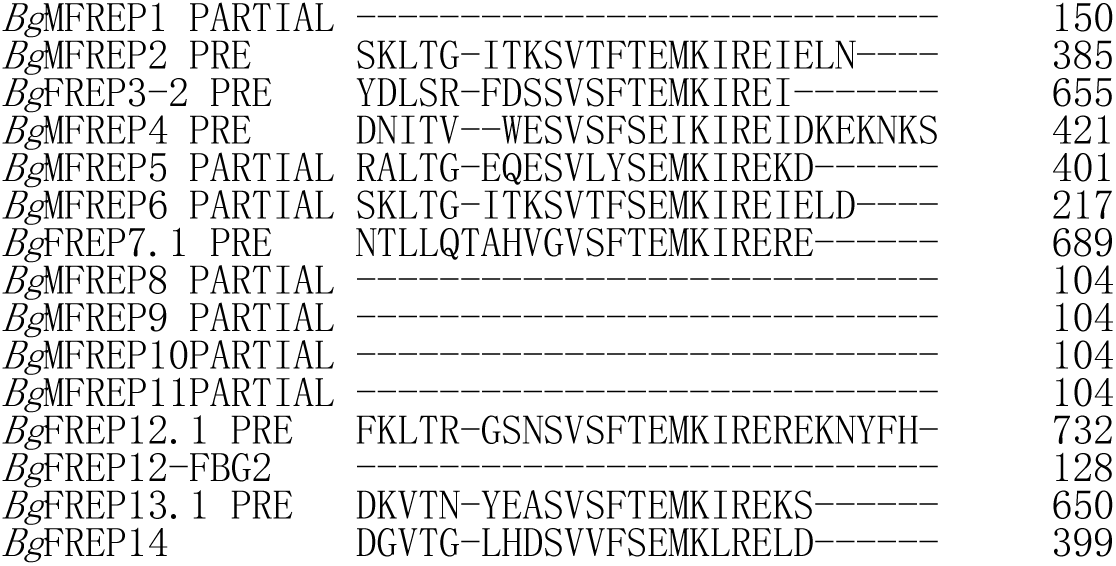
Alignment of multiple *Bg*FREP amino acid sequences and distribution of identified peptides. Peptides identified by LC-MS/MS are highlighted in gray. The GenBank accession numbers of each entry are: *Bg*MFREP1 partial, AAK13549; *Bg*MFREP2 precursor, AAK13550; *Bg*FREP3-2 precursor, AAK28656; *Bg*MFREP4 precursor, AAK13551; *Bg*MFREP5 partial, AAK13546; *Bg*MFREP6 partial, AAK13552; *Bg*FREP7.1 precursor, AAK28657; *Bg*MFREP8 partial, AAK13553; *Bg*MFREP9 partial, AAK13554; *Bg*MFREP10 partial, AAK13555; *Bg*MFREP11 partial, AAK13556; *Bg*FREP12.1 precursor, AAO59918; *Bg*FREP12-FBG2, AAT58639; *Bg*FREP13.1 precursor, AAO59922 and *Bg*FREP14, ABO61860.

**Figure 1—figure supplement 6.**
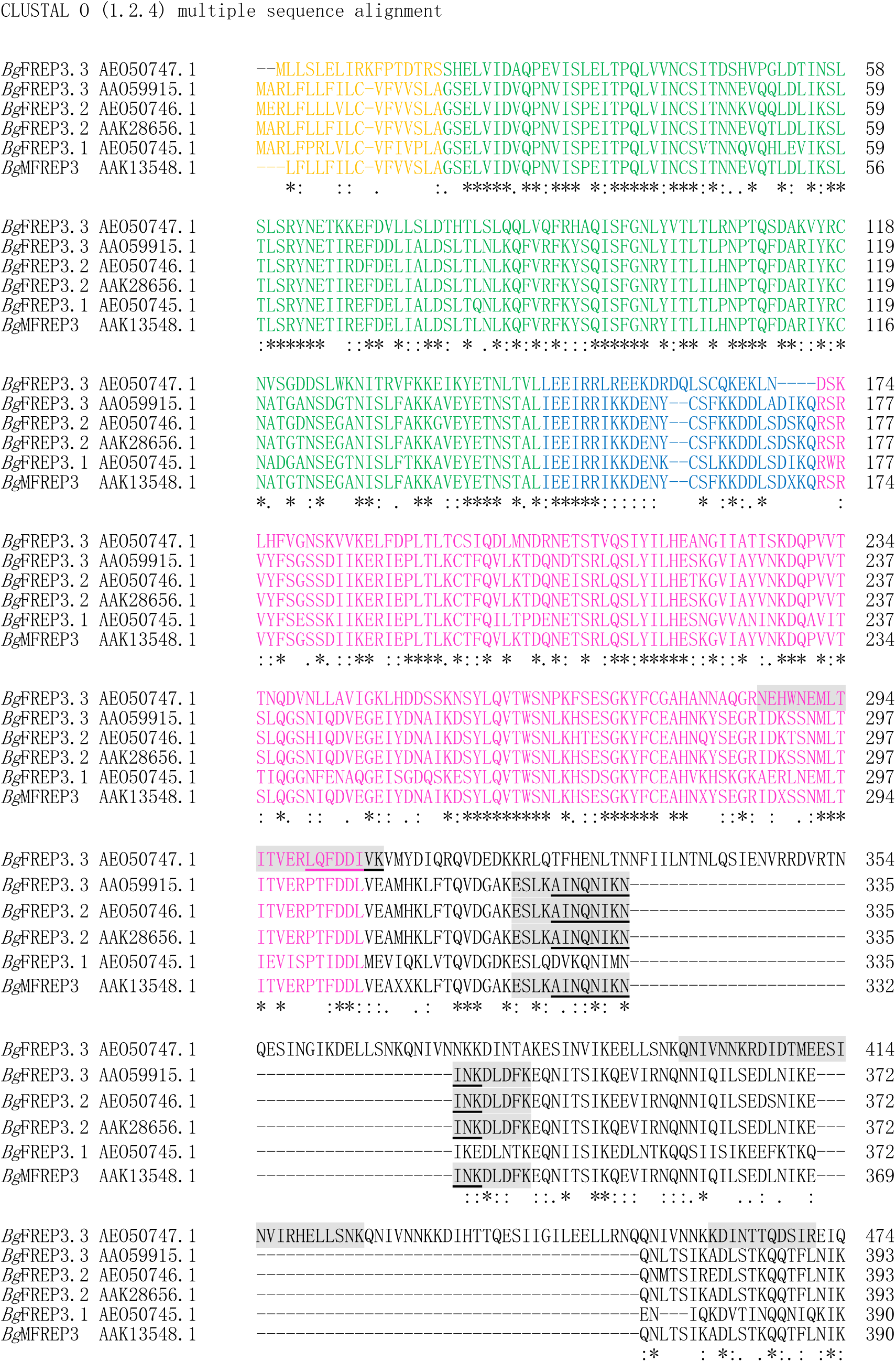
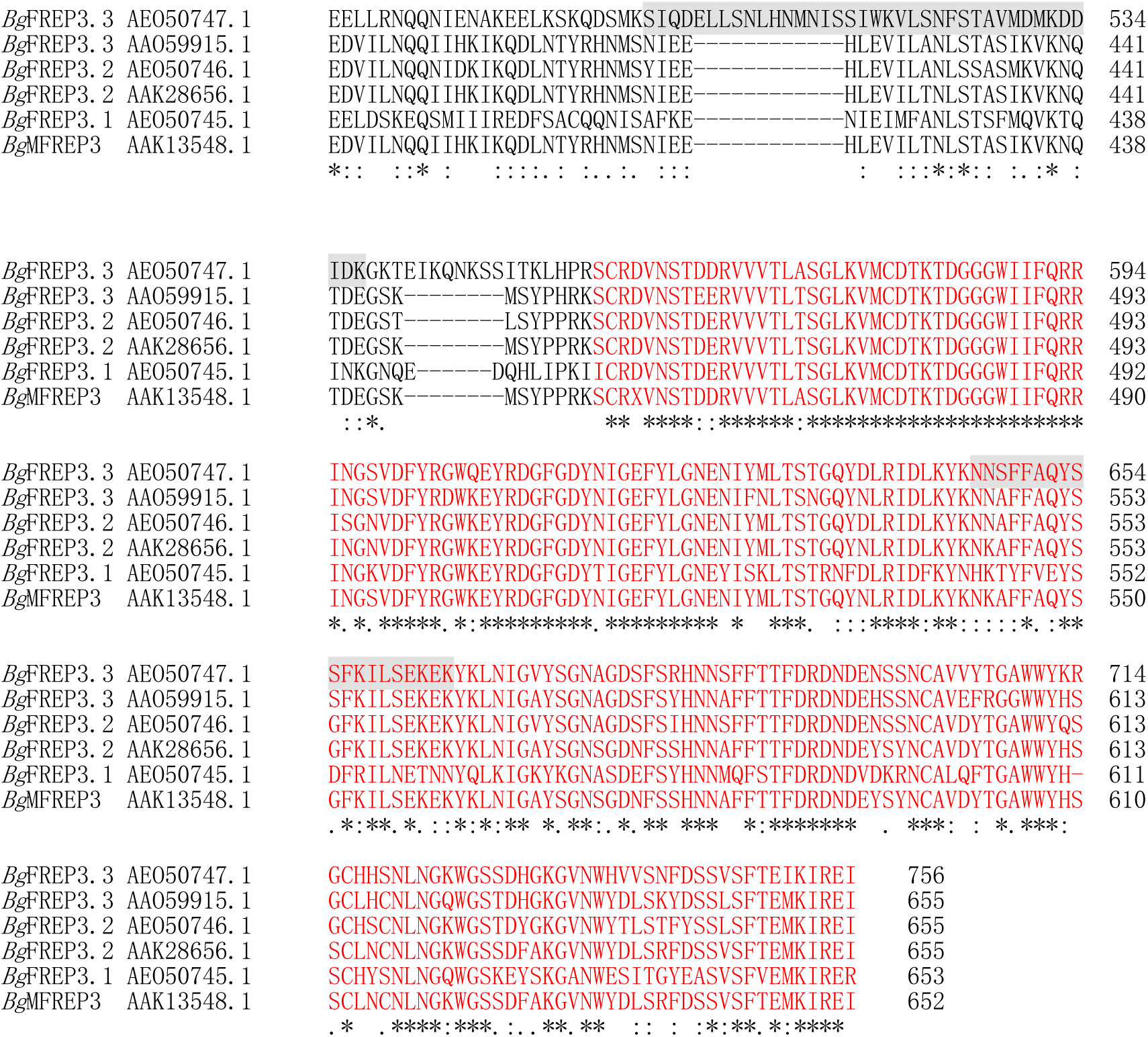
Alignment of multiple *Bg*FREP3 amino acid sequences and distribution of identified peptides. Peptides identified by LC-MS/MS are highlighted in gray, the overlapping region of peptides are bold underline. GenBank annotations of *Bg*FREP3.2 (AAK28656.1) are used for locate the individual *Bg*FREP3 domains: orange, signal peptide; green, IgSF1 domain; blue, small connecting region; pink, IgSF2 domain; black, ICR; red, FBG domain.

**Figure 4—figure supplement 1.**
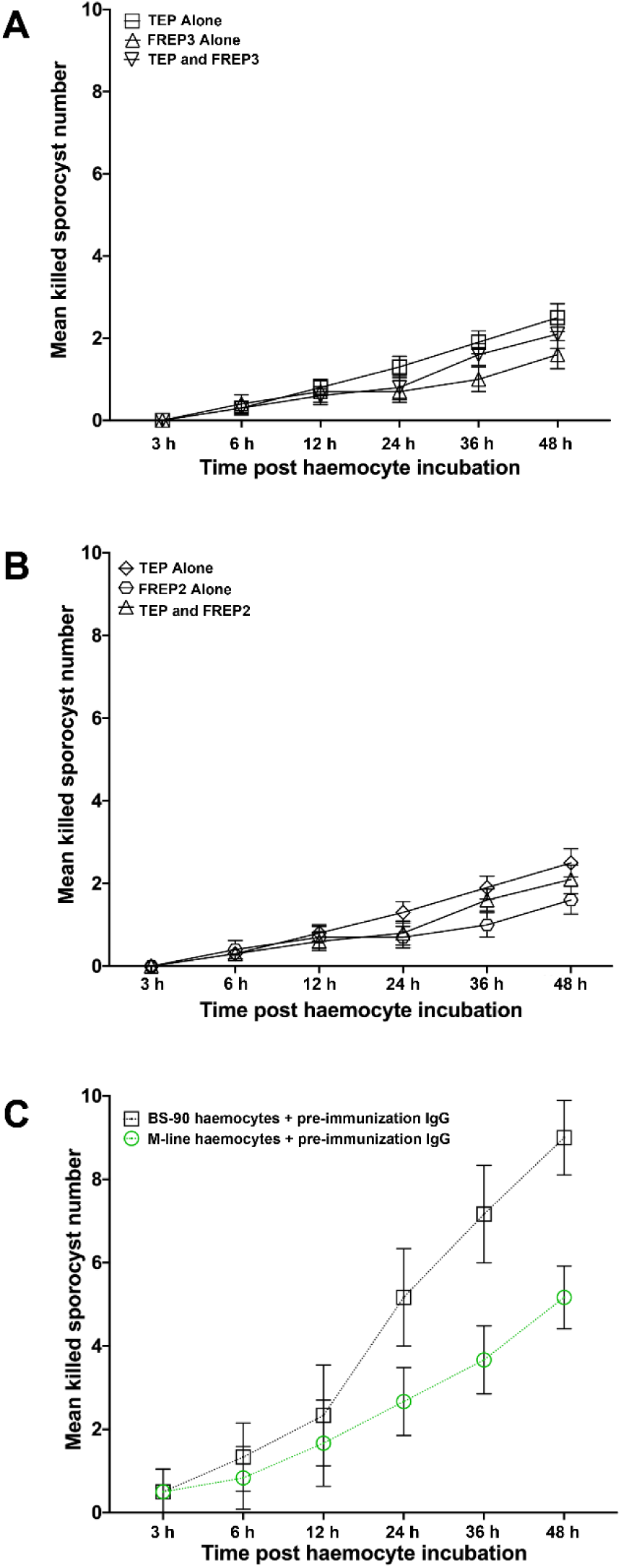
Controls of functional verification experiments. **A:** Only the media and recombinant proteins r*Bg*FREP3 and r*Bg*TEP1 were present, without snail plasma or haemocytes, to control the sporocyst killing assay of r*Bg*FREP3, r*Bg*TEP1 and the combination of both. **B:** The controls of the sporocyst killing assay of r*Bg*FREP2, r*Bg*TEP1 and r*Bg*FREP2-r*Bg*TEP1 complex. **C:** Pre-immune serum from the same rabbit which raised the anti-r*Bg*FREP3 antibodies were incubated with BS-90 and M-line haemocytes for control.

**Table S1.**
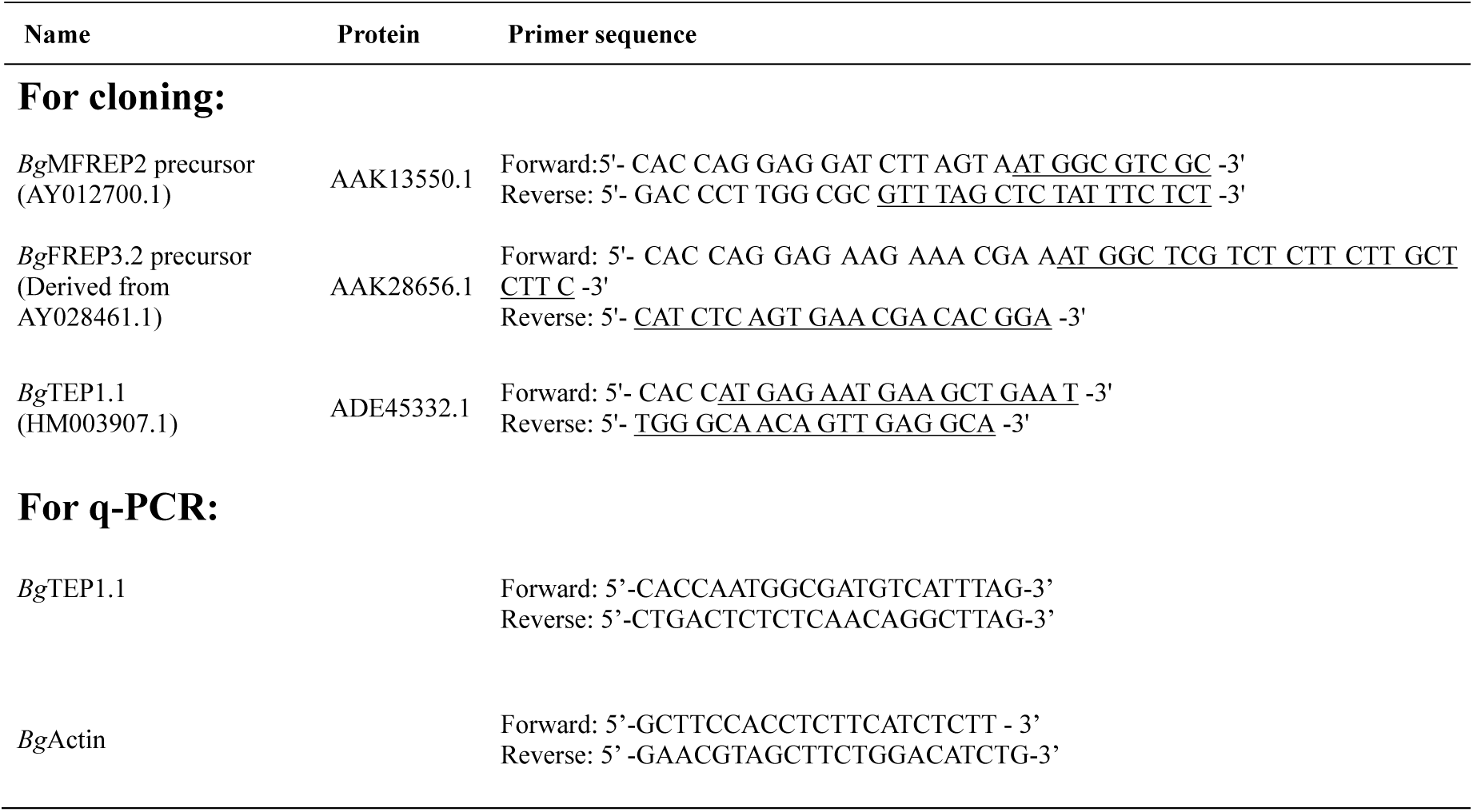
Primer list for cloning and quantitative RT-PCR.

**Table S2.**
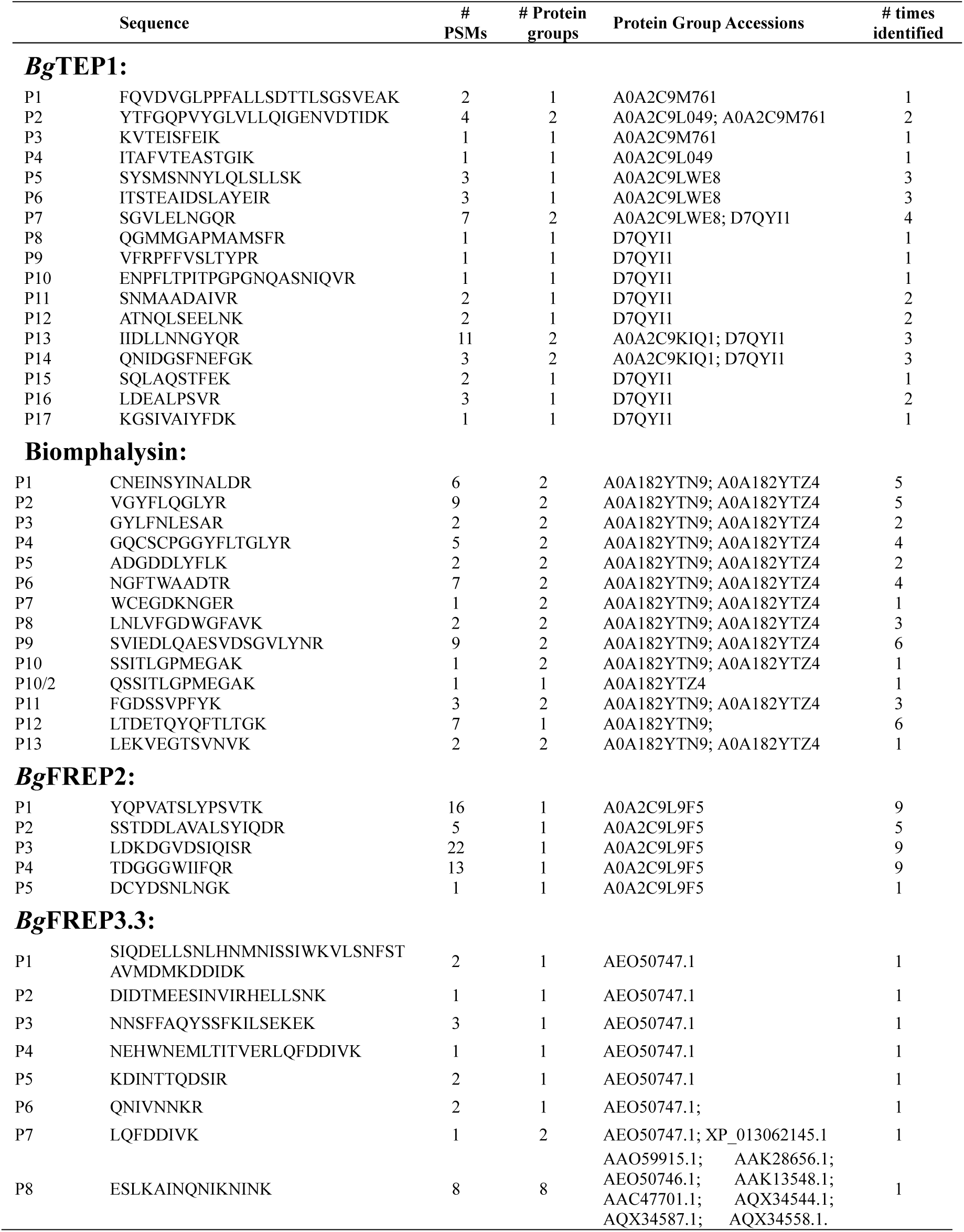
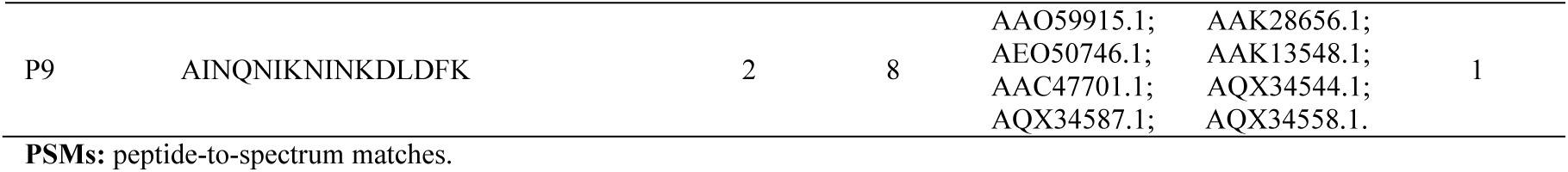
The identified peptides of *Bg*TEP1, Biomphalysin, *Bg*FREP2 and *Bg*FREP3.3 by LC-MS/MS.

